# DLC2Action: A Multimodal Deep Learning-based Toolbox for Automated Behavior Segmentation

**DOI:** 10.1101/2025.09.27.678941

**Authors:** Elizaveta Kozlova, Andy Bonnetto, Alexander Mathis

## Abstract

Behavioral analysis is fundamental to neuroscience, yet the manual annotation of actions remains a bottleneck that constrains both the scale and the reproducibility of experiment. Here, we present DLC2Action, an open-source Python toolbox that enables automatic behavior annotation from video, audio and estimated 2D/3D pose tracking data. DLC2Action integrates multiple state-of-the-art deep learning architectures optimized for action segmentation and supports self-supervised learning (SSL) to address annotation scarcity, boosting performance with limited labeled datasets. To enable model comparison, we establish fixed train/test partitions for nine diverse datasets comprising rodent experiments, human cooking studies, and wildlife observation. DLC2Action reached strong performance across those benchmarks. To further showcase the tool’s versatility, we applied it to Atari gameplay data and found that, in certain games, the players’ eye movements consistently predict their button presses across subjects. Because DLC2Action features an intuitive graphical user interface (GUI), users can streamline behavior annotation, perform active learning, and assess of model predictions. Diverse pose, video, and annotation formats are supported. Lastly, DLC2Action is modular and thus designed for extensibility, allowing users to integrate new models, dataset features, and methods. The code and benchmarks are available at: https://github.com/amathislab/DLC2action

## Introduction

Quantifying behavior is essential for psychology, ethology, medicine, neuroscience, and many other fields (1–6). For instance, in neuroscience, quantifying behavior is essential to characterize the role of neural circuits (7) or to optimize translational approaches (8). Manual behavior annotation can be precise but expensive in terms of time and human resources (3, 9). One way to tackle this problem is via unsupervised approaches, which do not require human annotations a priori. However, while many excellent tools (10–19) have emerged, their output must still be validated by experts, which often requires as much interpretive effort as a priori annotations.

Teaching machines how to identify behaviors in a supervised way, that is, based on expert annotations, is challenging. Distinct behaviors can appear visually similar (e.g., resting and freezing in mice). Conversely, the same behavior can occur in many contexts with highly varying appearances (e.g., reaching for a cup of espresso in a dark Manhattan bar vs. a bright Neapolitan café). This problem can be mitigated by using a more appropriate representation. For instance, reaching can be nicely predicted from 3D pose estimation data, irrespective of luminance. In contrast, freezing vs. resting could be better predicted from fur texture changes due to physiological changes rather than from posture. Kinematic features capture what the body does rather than how it looks, making them robust to lighting variation, background clutter, and camera angle in ways that raw pixel representations are not. This is why tools should operate on pose-derived features by default while retaining the option to incorporate video and audio when audiovisual context is informative.

Here, we introduce DLC2Action, a novel toolbox for performing action segmentation based on pose and video features. Indeed, kinematic information relevant for behavioral analysis can be efficiently extracted with little keypoint annotation data based on pose estimation (20–26) or high-level video features extracted from encoding models such as VideoMAE or DINOv3 (27), which were pretrained on large-scale video datasets (28–30). Based on both of these features, one can train relatively lightweight architectures to perform action segmentation, i.e., the task of predicting what behavior is being carried out in each frame of a video. We want to emphasize that DLC2Action includes general-purpose sequence-to-sequence models and can, in principle, deal with more modalities. We illustrate this flexibility by predicting gameplay from eye-tracking data during video gameplay in humans.

Multiple action segmentation tools exist and are used in Neuroscience, such as JAABA (31), ETH-DLCanalyzer (32), MARS (23), DeepOF (33), SimBA (34), and A-SOiD (35).

There are a number of differences in comparison to these tools, but most prominently, DLC2Action is more flexible, allows multimodal input, standardizes several benchmarks for future model development, and provides easy access to multiple state-of-the-art methods from the action segmentation literature, which we have also adapted to improve performance. DLC2Action addresses these gaps by offering a common interface for multiple state-of-the-art architectures, working with any pose estimation output, and — critically — providing standardized benchmarks that none of the above tools currently supply.

DLC2Action brings together eight temporal deep learning architectures — including multi-stage temporal convolutional networks (TCNs) and attention-based transformers — that achieve strong performance in action segmentation benchmarks in computer vision (36–42) and adapts them to operate on multimodal features rather than raw video. To address the absence of community-standard evaluation protocols in behavioral neuroscience, we fix train/test splits for nine datasets and report baseline performance for all architectures (Table 1), creating reference points against which future tools can be directly compared. The toolbox is modular, allowing users to add new models, losses, and data formats without modifying the core pipeline.

**Table 1.** Overview over benchmarking datasets: **Split types**: Time:strict methods strictly separate the training from the validation set and avoid overlaps (see Methods). File methods select different video-annotation pairs for the training and testing set. **Orig. split**: indicates whether the original split is used (Yes) or a new division is proposed (No). **Leading model**: Leading model with respect to the split. *hBehaveMAE only uses self-supervised methods. **The authors used DLC2Action.

| Dataset | Split type | Orig. split | Leading model |
| --- | --- | --- | --- |
| EPM (32) | Time:strict | No | N/A |
| OFT (32) | Leave-one-out | Yes | SIPEC (48) |
| SimBA-RAT (34) | Time:strict | No | N/A |
| CRIM13 (47) | Time:strict | No | N/A |
| CalMS21 (45) | File | Yes | ResNet/CPC (49) |
| Shot7M2 (50) | File | Yes | hBehavMAE* (50) |
| hBABEL (50) | File | Yes | hBehavMAE* (50) |
| ESK (43) | File | Yes | C2F-TCN** |
| MammAlps (46) | File | No | N/A |

## Results

Fundamentally, DLC2Action solves action segmentation problems, i.e., predicting what behavior is present on a frame-by-frame basis, and it can do so from either pose estimation representations, audio, or video features (Figure 1A). Although DLC2Action’s name suggests that it takes DeepLabCut’s output (DLC (20, 22, 24)) to predict actions, it is versatile and can be used in combination with other pose estimation packages, directly with manually extracted features, or for different behavior annotation data formats. In essence, DLC2Action is a Python package that provides a high-level API to perform behavioral analysis with a graphical user interface (GUI). Here, we describe its workflow, interface, and features. The description stands for version 1.0, the code is available at https://github.com/amathislab/DLC2action.

**Figure 1.**
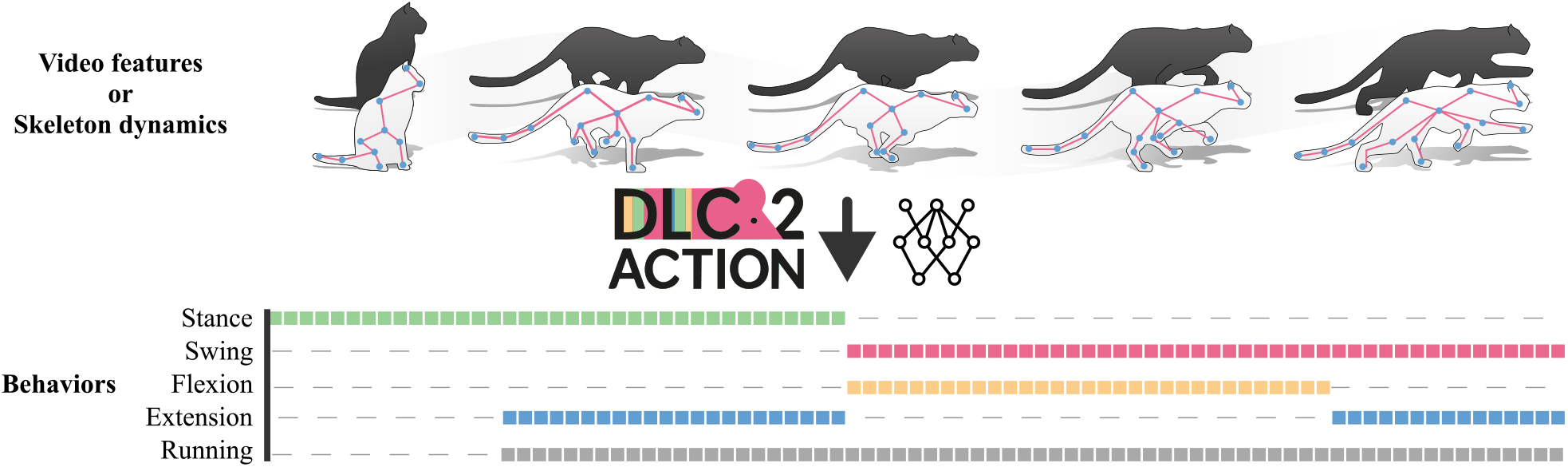
Overview of DLC2Action. DLC2Action takes pose estimation data or video features as input and uses state-of-the-art neural networks to predict behaviors. This architecture-agnostic design means the same pipeline applies to a mouse in a maze, a human in a kitchen, or a chamois in the Swiss National Park.

### Workflow and overview

Reliable behavior labels are essential for downstream analysis, but getting from raw pose tracks to those labels is typically an iterative, multi-step process; DLC2Action is designed to support this process. In practice, this process spans dataset loading and preprocessing, feature extraction, dataset splitting, model training and hyperparameter tuning, prediction review and correction, and finally deployment to annotate new videos at scale.

DLC2Action covers the full annotation workflow — from importing existing labels or creating new ones via the GUI, through training and hyperparameter search, to reviewing predictions (proofreading) and deploying the best model at scale — within a single project environment (Figure S1A, full workflow in Methods).

At its core, DLC2Action provides an accessible yet comprehensive deep-learning training pipeline. Starting from single or multi-agent skeleton dynamics (Figure 2A), the system extracts kinematic features and learns to map them onto user-defined behavioral categories. Eight model architectures are available (Figure 2B), comprising temporal convolutional networks and transformer architectures. They can be trained individually or in combination to maximize performance. Self-supervised learning modules can further boost performance when labeled data are scarce. During training, data are divided into fixed-length, potentially overlapping segments to increase pattern redundancy and improve generalization.

**Figure 2.**
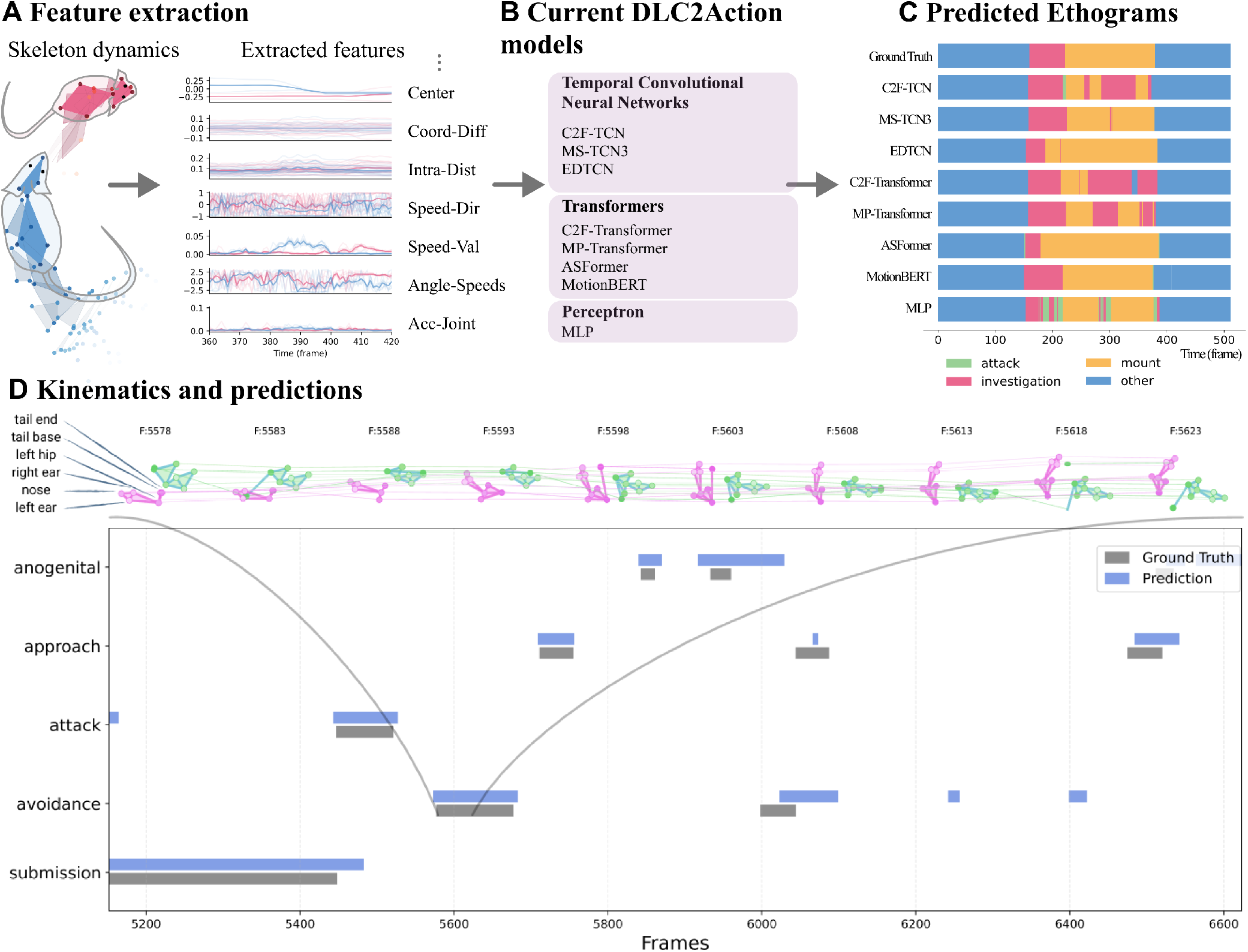
Architecture details and worked example for the CalMS21 dataset (45) and SimBA-RAT (34). (A) Feature extraction. From skeleton dynamics, DLC2Action extracts several kinematic features including joint accelerations. (C) Predicted Ethograms. DLC2Action proposes a set of functions to plot performance curves and ethograms of the models predictions. (B) DLC2Action models currently implemented in the package include temporal convolutional neural networks, transformers and one multi-layer perceptron model. (D) Kinematics and predictions for the SimBA-RAT dataset. The kinematics (top) are sampled from the first 45 frames of an “avoidance” segment, where the purple mouse moves around the green mouse. The predictions (bottom) are from the C2F-TCN model and show good alignment with the ground-truth annotations.

Three of the eight architectures were originally designed for video-based action segmentation and were adapted here to accept kinematic features as input (Figure S1). Diagnostic outputs such as loss curves, example predictions (Figure 2C), and segmentation statistics can be generated to help users assess and iterate on model performance (Figure S11). Together, these features establish a robust workflow for connecting kinematic data to complex behaviors (Figure 2D).

### Interfaces

DLC2Action provides a graphical-user interface (GUI), as we will detail later. Furthermore, DLC2Action provides a high-level Application Programming Interface (API) to interact with the data. Only four lines of code (Code block 1) are sufficient to go from project creation to inference. The API documentation is available online.

### Multimodal Action segmentation

Intuitively, some behaviors can be better predicted from video rather than 2D/3D pose estimation. DLC2Action includes the possibility of using external features in addition to kinematic information, such features coming from videos or audio (Figure 3A). To do this, we concatenate the different features before feeding them to DLC2Action models. We assessed the performance of the DLC2Action models when leveraging videos by extracting representations from the powerful video model VideoMAE (30), which was pre-trained on the egocentric view of the EPFL-Smart-Kitchen-30 dataset (43) (Figure 3B). By concatenating the 3D body and hand pose information with (learned) visual features, we can achieve the best performance in activity segmentation (Figure 3C and detailed results in (43)). Conversely, leveraging powerful video encoding only slightly enhances the performance of DLC2Action on the Open Field Test dataset (OFT) (32) (Figure 3B and Figure S9), where models based on kinematic features have already reached a strong performance of 64.0% F1. Here we used a VideoMAE model that was pre-trained on the large video action recognition dataset Kinetics-700 (44).

**Figure 3.**
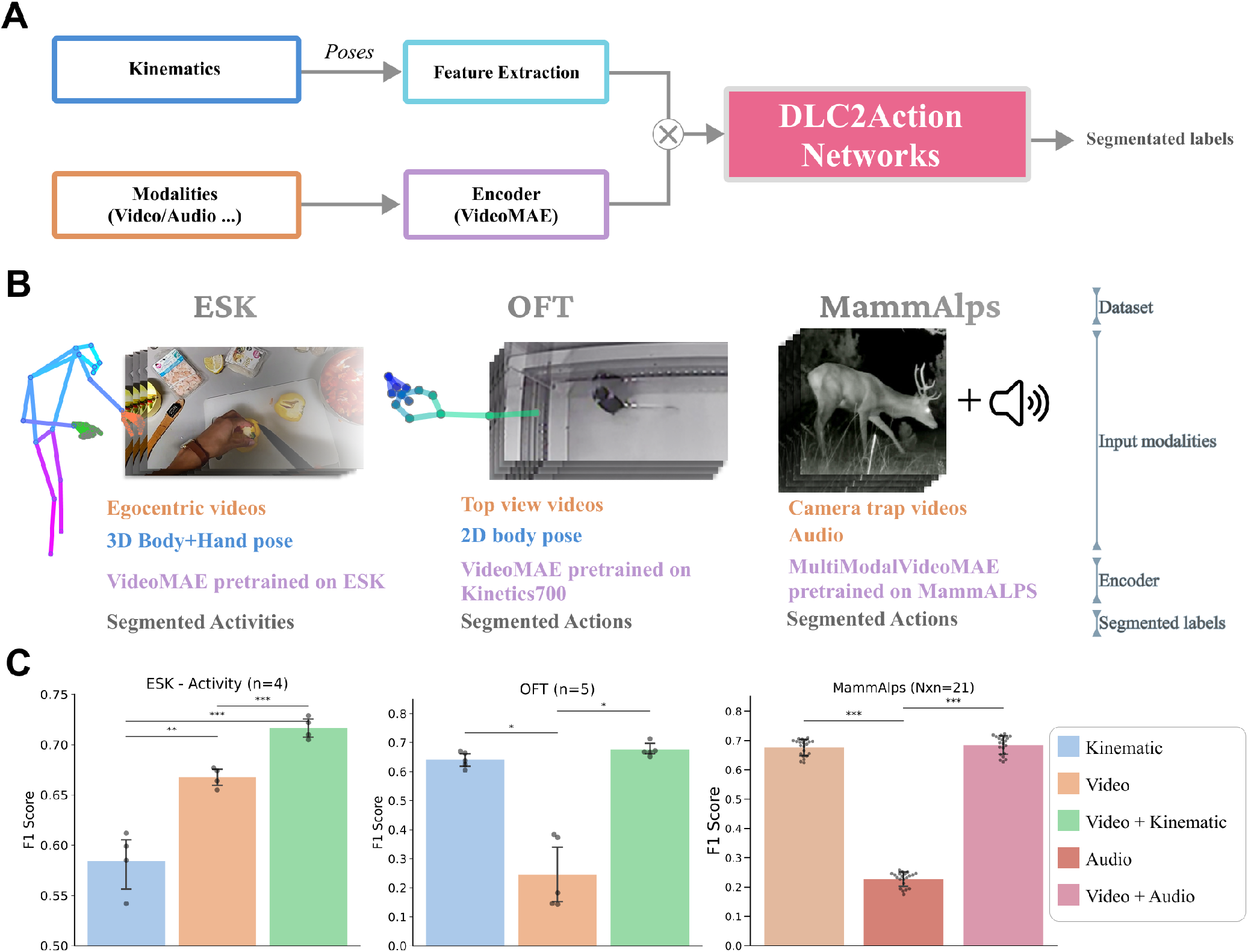
Multimodality in DLC2Action. (A) Multimodal fusion architecture, kinematic information are concatenated with deep video or/and deep audio features from a VideoMAE model (30) pretrained on the specific modalities before being fed to DLC2Action. (B) Configuration of experiments for the ESK dataset (43) (left), the Open Field Test (OFT) (32) dataset (middle), and the MammAlps dataset (46) (right). ESK and OFT combine videos kinematic representations, while MammAlps compare video and audio modalities. (C) Performance of the DLC2Action model with multimodal inputs. For the ESK dataset (left), video features provide a significant contribution to prediction F1 score. In contrast, for the OFT dataset (middle), kinematic information alone is sufficient to reach high performance, with minimal additional benefit from video features. For the MammAlps dataset (right), adding audio to video input only slightly improve the performance. Error bars correspond to standard deviation over different model architectures (*m*) or different splits and model architectures (*m × n*). Statistical significance was assessed using a one-way ANOVA with post-hoc *t*-tests for the OFT dataset (normal distribution), and a Kruskal–Wallis test with post-hoc pairwise Mann–Whitney *U* tests for the ESK and MammAlps datasets (non-normal distribution). Asterisks denote statistically significant differences between the indicated groups (\**p <* 0.05, \*\**p <* 0.01, \*\*\**p <* 0.001). Comparisons unmarked were not statistically significant (*p >* 0.05).

**Code block 1.**
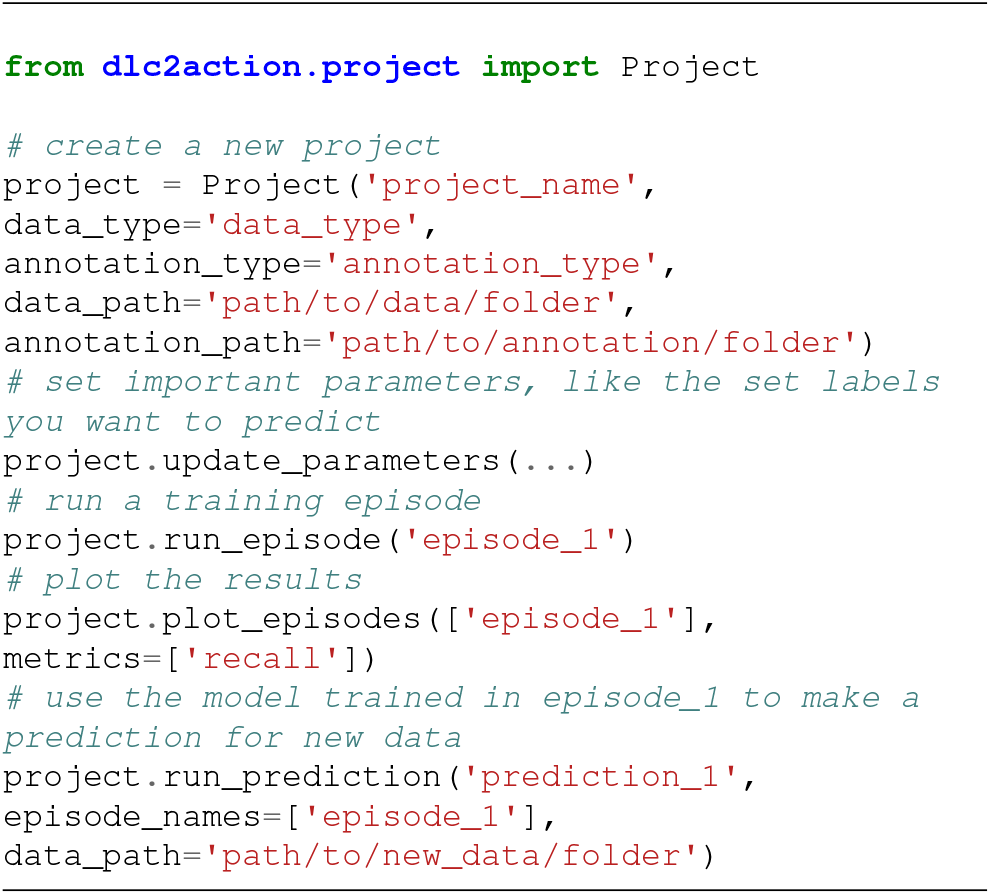
Minimal API calls for creating and configuring a project, training a deep learning model and performing action segmentation on a folder.

The ability to leverage multiple modalities in DLC2action is flexible. If one can extract features from the modalities of interest, DLC2Action can then handle action segmentation. In MammAlps (46), the authors collected camera-trap videos and audio of different species observed in the wild. Using an adapted VideoMAE model that accepts both video and audio (via spectrogram conversion) and is fine-tuned on the dataset. We observe a performance boost when using video with audio features compared with video only (Figure 3C, and Figure S6A). While the (average) performance when using video with and without audio is similar, some behaviors, such as jumping and scratching antlers, are much better predicted (Figure S6B); indeed, when we consider behaviors with the highest sound intensity, including audio, boosts performance (Figure S7). See Methods for details on how the dataset was converted into a (novel) action-segmentation benchmark.

Overall, this illustrates that DLC2Action can leverage multiple modalities for action segmentation. To rigorously evaluate DLC2Action, we curated nine public action segmentation datasets. These diverse datasets include individual and social behaviors, featured both humans and animals, and encompassed exclusive and multi-label classification tasks.

### Benchmarking across animal and human datasets

The datasets were sourced from synthetic, lab-controlled, and real-world environments. For all of them (except MammAlps (46)), a set of kinematic features was extracted from the available pose estimation data. When no public split was available, we split the data into training and test sets. We fixed splits for four action segmentation datasets, namely EPM (32), CRIM13 (47), SimBA-RAT (34) and MammAlps (46). For EPM, SimBA-RAT, and CRIM13, we employed “Time:strict” splits to test temporal generalization—how well models predict behavior at unseen time points within individual recordings. For MammAlps, we created file-based splits to assess generalization across multiple individuals and ecological contexts. For OFT and CalMS21, we followed established protocols to evaluate generalization to novel individuals. For Shot7M2, hBABEL, and ESK, we used file-based splits to test generalization across recording instances and contexts. Overall, this diversity of evaluation strategies reflects different generalization goals relevant for different users. To support future benchmarking and reproducibility, we encourage the community to use these standardized train/test splits. Further details on the experiments and the train/test splits can be found in the methods (Table 1).

### Single-individual datasets

We first tested DLC2Action on datasets comprising widely used behavioral tests in rodents, namely the elevated plus maze (EPM) (Figure 4A) and the OFT (Figure 4B), generated by Sturman et al. (32). For the OFT dataset, *grooming, supported rearing*, and *unsupported rearing* were annotated, while *grooming, head dip, protected stretch*, and *rearing* were annotated in the EPM dataset. The mice were tracked with DeepLabCut (20) (see Methods). We did not find any baseline methods evaluating the EPM dataset; therefore, we propose a time-based split and turn this dataset into a benchmark (see Methods). MotionBERT (42) yields the best performance on EPM with an F1 score of 70.0% ± 7.44% (standard deviation calculated over splits). On the OFT dataset, we found that the DLC2Action models outperform the multilayer perceptron reported in the original study (32) as well as a classifier that additionally uses visual features (48). As in the original studies, the results were obtained using a leave-one-out cross-validation method (N = 20 videos). In Sturman et al. (32), the average F1 score is 57% ± 13%; in Marks et al. (48) it is 64% ± 15%, while our C2F-TCN model achieves 74.5% ± 11.4% without visual features (standard deviation calculated over splits).

**Figure 4.**
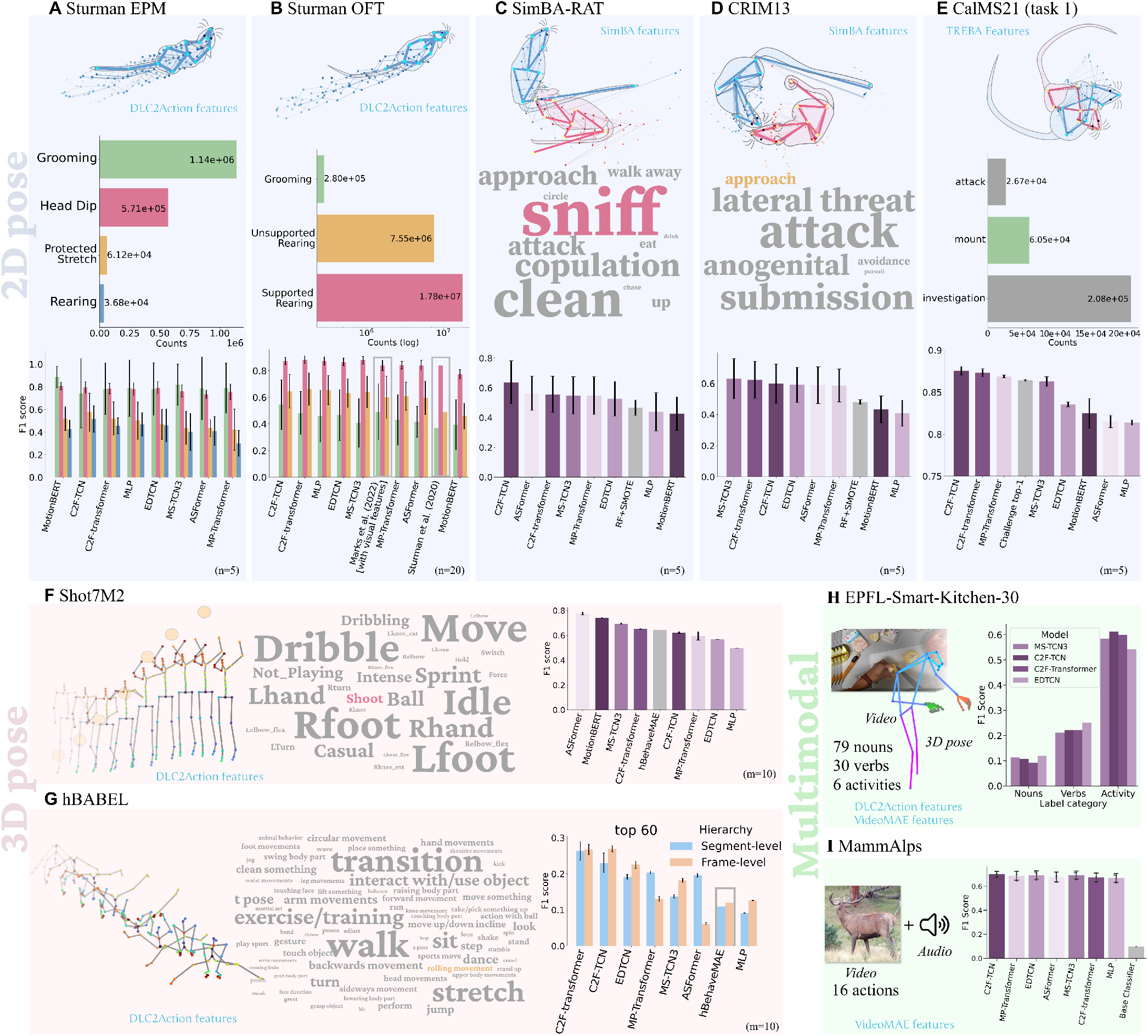
Comprehensive Benchmarking Suite. Datasets are illustrated by an example pose with keypoints from previous timepoints, the distribution of labels and the F1 score performance of each model. We also report the type of features used as input to the models (cyan). The word clouds illustrate the different actions, with the word sizes proportional to the action frequency. We report the performance on rodent 2D pose estimation data (A-E) and human 3D pose estimation data (F-G) and multimodal data (H-I). Performance is reported per class (A-B) or averaged over class (C-G). Baselines are represented in gray (filled bar (C-I) or contours (A,G)). We applied DLC2Action to (A) Elevated platform maze (EPM) (n=5) (B) Open Field Test (OFT) (n=20). Baseline results from (32) and (48). (C) SimBA RAT (n=5). RT+SMOTE results are used as baseline (see Methods). (D) CRIM13 (n=5). RT+SMOTE results are used as baseline (see Methods) (E) CalMS21 task-1 (m=5). Challenge top-1 results from (45) (F) Shot7M2 (m=10, MotionBERT m=2). Linear probing results with hBehaveMAE from (19). (G) hBABEL (m=10, MotionBERT m=2) results for different models. Linear probing results with hBehaveMAE from (19). (H) EPFL-Smart-Kitchen-30 (m=1). Multimodal dataset with video and pose features as input. DLC2Action was used as the baseline of the original publication (43) (I) MammAlps (n=3). Multimodal dataset with video and audio features as input (46) Across the benchmark suite, DLC2Action matches or exceeds prior methods. The error is reported over different train/test splits (m) for each model or different initialization seeds (n).

### Multi-individual datasets

In order to show the flexibility of DLC2Action, we evaluated our framework on social behaviors in rodents based on 2D pose estimation data. In particular, we used the datasets presented in SimBA (34), comprising their rat resident-intruder dataset (SimBA-RAT) and the pose estimation versions of CRIM13 (47) as well as CalMS21 (45).

The CRIM13 dataset contains manual annotations for *anogenital sniffing, approach, attack, avoidance, boxing, lateral threat*, and *submission* in mice, while SimBA-RAT involves a mix of individual and interactive behaviors in rats (*copulation, sniffing, drinking*, etc.) (Figure 4C-D). We compare the performance of DLC2Action on this data to that of a random forest model (51) in combination with the SMOTE (52) resampling method. Our models perform better here as well (63.8 *±* 14.5% vs 46.7 *±* 5.2% for the SimBA-RAT dataset with the C2F-TCN model, and 63.1 *±* 12.9% vs 48.2 *±* 1.5% for the CRIM13 dataset).

From the CalMS21 benchmarking challenge (45), we focus on the classic action annotation scenario. The task involves the segmentation of three social behaviors (attack, mounting, and close investigation) (Figure 4E). The training data is composed of 70 videos that last from 3 seconds to 12 minutes, and the test data consists of videos annotated by the same expert. DLC2Action models slightly outperform the top-1 model from the challenge (87.6 ± 0.5% F1 score for the best DLC2Action model vs. 86.4 ± 0.1% for the top-1 model in the challenge) (standard deviation over 5 model initializations). Note that while the existing top-1 model was trained using large unannotated datasets, this was not needed for DLC2Action models to match their performance.

### Action segmentation for 3D pose estimation data

To benchmark the performance of DLC2Action with 3D pose data, we tested the models on the Shot7M2 and hBABEL datasets (19) (Figure 4G), as well as on the EPFL Smart kitchen benchmark (43) (Figure 4H). These benchmarks are compared with hBehaveMAE (19), an unsupervised behavior representation method that is the current state-of-the-art representation learning approach in these benchmarks.

These datasets are pose-based hierarchical action segmentation benchmarks and provide dense, non-exclusive action annotations. These datasets are particularly challenging due to their large number of classes.

Shot7M2 is a synthetic dataset composed of 7.2M fully annotated frames, featuring a 3D basketball player performing twelve actions and 14 movements (short-term behaviors) across four different activities (long-term behaviors). For example, during an *intense playing* activity, the player can perform actions such as *sprint, dribbling* and *shoot*, while simultaneously annotated movemes include *elbow flexion, knee extension, left foot* ground contact, and *ball*-hand contact. DLC2Action models achieve strong performance on this benchmark, outperforming both a simple MLP and existing methods (Figure 4F). On Shot7M2, the best DLC2Action model achieves 77.6 *±* 1.1% F1 compared to 64.7% for hBehaveMAE (50), a self-supervised method with linear probing. This performance difference is expected, as linear probing (which freezes pretrained representations) is substantially weaker than the end-to-end fine-tuning used in DLC2Action.

hBABEL is an extension of the BABEL dataset (54), transforming it into a non-exclusive hierarchical action segmentation dataset. It comprises 6.2M frames and overlaps long-term and short-term behaviors with over 180 different classes (selected from the most frequent behaviors to ensure a similar distribution of labels in the training and testing sets). The 3D position of 25 major joints was processed from the AMASS dataset, which was collected using motion capture (55). Despite the challenge posed by the large number of classes, DLC2Action substantially outperforms both a simple MLP and the SSL baseline hBehaveMAE (50), achieving roughly twice the performance of the latter: 26.4% ± 2.7% vs 10.9% (segment-level) and 26.8% ± 1.4% vs 12.0% (frame-level) F1 scores.

The EPFL-Smart-Kitchen-30 dataset (ESK) (43) contains 29.7 hours of densely annotated cooking behaviors, totaling 60,189 segments at an average rate of 33.78 segments per minute for 763 fine-grained actions that can be decomposed into 33 verbs, 79 nouns, and 5 activities corresponding to longer behaviors. Using video data collected from nine exocentric cameras and one egocentric camera, the authors inferred 3D body and hand pose information. DLC2Action achieves 35.0% F1 on verb segmentation with the C2F-Transformer model, 35.2% F1 on nouns with the C2F-TCN model, and 72.9% F1 on activity prediction with the MS-TCN3 model using combined video features and kinematic information as input.

### Action segmentation in the wild

The MammAlps dataset (46) is composed of 8.5 hours of densely annotated animal recordings collected from camera traps placed in the Swiss National Park. The dataset includes individual tracks with both the species and the behaviors annotated. In our case, we turned the data into an action segmentation problem (see Methods) and considered 15 frequent behaviors, including *grazing, sniffing*, and *standing head up*. The extracted dataset is composed of 4,206 segments, which are split into clips-based on a 70%/30% train/test split (Figure S8). Using both video and audio data, we obtain 70% F1 (Figures 4I and S5).

### Predicting gameplay from saccades in humans playing video games

Behavioral tasks require the coordination of different body parts (56). More specifically, when playing video games, certain games will likely require the coordination of eye movements and actions. While one can likely just overfit to the idiosyncratic behavior of particular players, we consider the more interesting case. Namely, if common eye-action coordination emerges for particular games across individuals. Here we test this hypothesis based on the Atari-HEAD dataset with DLC2Action. The Atari-HEAD dataset boasts 117 hours of gameplay data from 20 different games, with eight million action demonstrations and 328 million gaze samples (53). To demonstrate the potential of DLC2Action for diverse applications, we tested whether we could predict game commands from the eye gaze trajectories without considering the screen content (Figure 5A).

**Figure 5.**
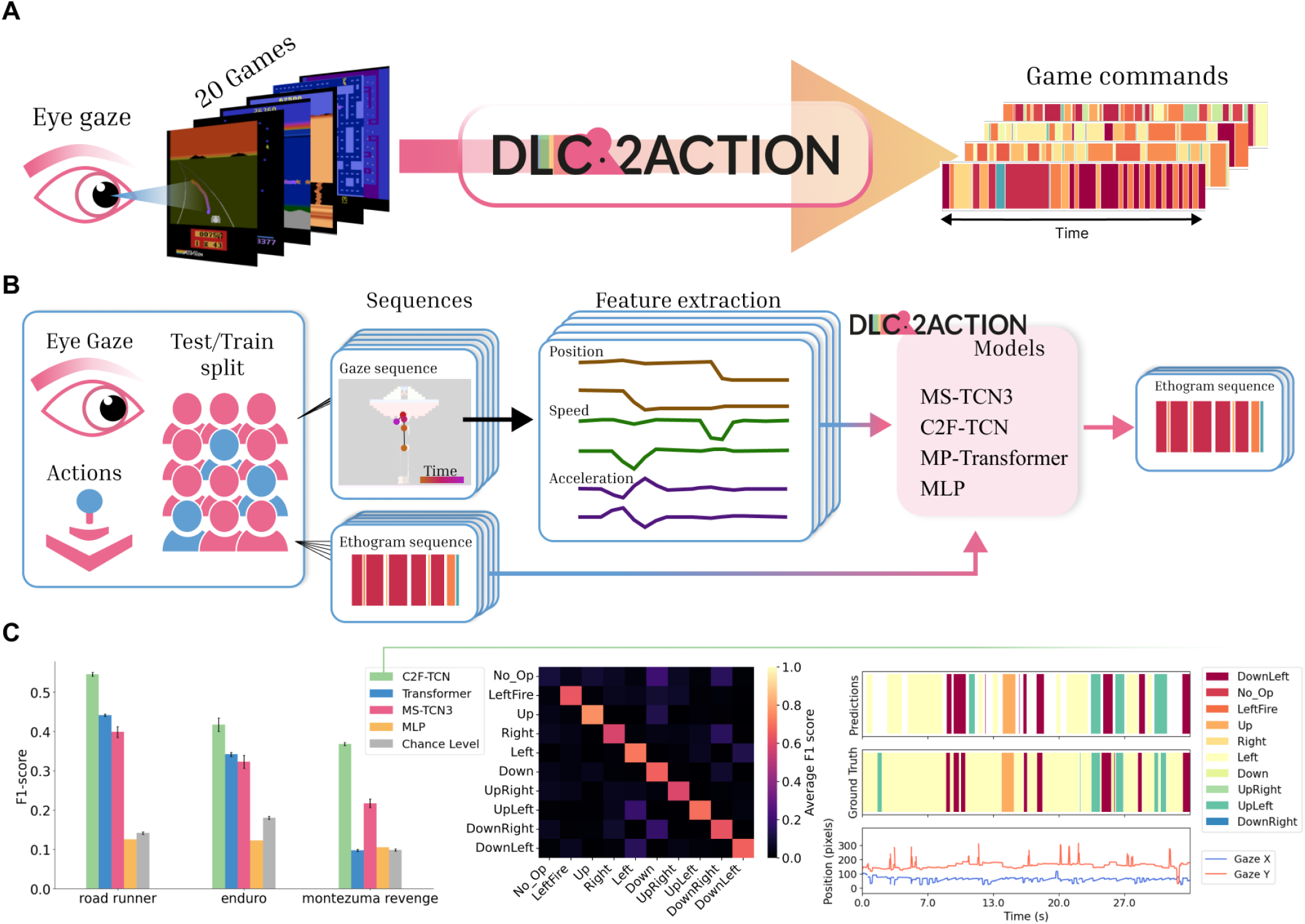
Prediction of Atari-HEAD video game commands from eye gaze. (A) Workflow. 17 game commands were predicted from 2D eye gaze movements among 20 video games played by 25 participants using DLC2Action. (B) Detailed workflow. The data was split with a 80/20 Train/Test split across participants. Pose, speed and acceleration were extracted and fed into 4 DLC2Action models to predict the actions. Chance level was also evaluated (see Methods). (C) Results. (Left) F1-score per model and per game for the three best performing games (Error bars: standard deviation across 5 splits) (Center) Confusion matrix for Road Runner using C2F-TCN. (Right) Predictions. (Top) Example of predictions in Road Runner when using C2F-TCN. (Middle) Ground truth game commands reference. (Bottom) Coordinates of the gaze in pixel units along the horizontal (X) and vertical (Y) axes. Experimental data from (53)

We compare the performance of four DLC2Action models (C2F-TCN, MS-TCN3, Transformer, and MLP) with a chance-level classifier that always predicts the most probable action over a whole episode, only considering the distribution of actions per video. For each game, we split the dataset into an 80-20% training-testing ratio, ensuring that entire videos (for distinct players) were present in the testing set (Figure 5B). The best performance was achieved for the game Road Runner with the C2F-TCN model (F1 = 54.53%), followed by the Enduro game and Montezuma Revenge (Figure 5C). These predictions have substantially better F1 scores than chance level. On the other hand, action commands from games such as Bank Heist or Riverraid could not be predicted reliably (Supplementary Figure S13). Thus, considering our choice of splitting methods, we found that some game-dependent decision strategies are not only correlated with gaze patterns but also consistent across participants for games with higher prediction scores. Notably, different temporal models achieved markedly different success rates on the same games—some models reaching above-chance F1 scores while others (such as MLP) performed near chance—suggesting that gaze-action relationships involve nonlinear or temporal structure that deeper temporal models can exploit. For the sake of this study, this brief analysis demonstrates both the flexibility of DLC2Action as an experimental framework to also study cognitive behaviors. Lastly, we will assess a few design decisions and describe the GUI in more detail.

### Kinematic features

Kinematic preprocessing is not a cosmetic step: replacing extracted features with raw keypoint coordinates consistently reduces F1 across datasets and architectures (Figure S S10), confirming that speed, acceleration, angular velocity, etc. (see Methods) carry behavioral information that absolute position alone does not encode. Users should test different combinations of kinematic features with respect to their behaviors of interest.

### Self-supervised learning experiments

While the automation of pose estimation can allow for measuring large amounts of kinematic data, the coverage of manual behavior annotations is usually lower. The introduction of a SSL module offers the possibility for the models to leverage both labeled and unlabeled datasets. DLC2Action supports three categories of SSL tasks, which can also be combined (Figure S12A-C and Methods).

We compared the performance of the C2F-TCN model both with and without the SSL module, which was also trained on the unlabeled data while using 5%, 10%, and 25% of the labeled data. On the OFT dataset (32), the SSL module can help boost performance with a low amount of labeled data and converges towards the reference performance when using 25% of the dataset as the training set (Figure S12D).

### Annotation interface

We also developed a graphical annotation interface that can be used in combination with the action segmentation package. The code is available at: https://github.com/amathislab/dlc2action_annotation

Upon visualizing selected video(s) and their respective pose estimation data, the annotation interface allows the user to interactively create simultaneous label classes in time segments using keyboard shortcuts. Editing tools allow the user to manipulate the labels or add uncertainty with single-frame precision (Figure 6A). The annotation interface supports multi-animal sequences and nested behavior annotations. It also supports displaying a wide range of additional data, including multiview cameras, pose skeletons, segmentation masks, and 3D keypoints with reprojection (Figure 6C).

**Figure 6.**
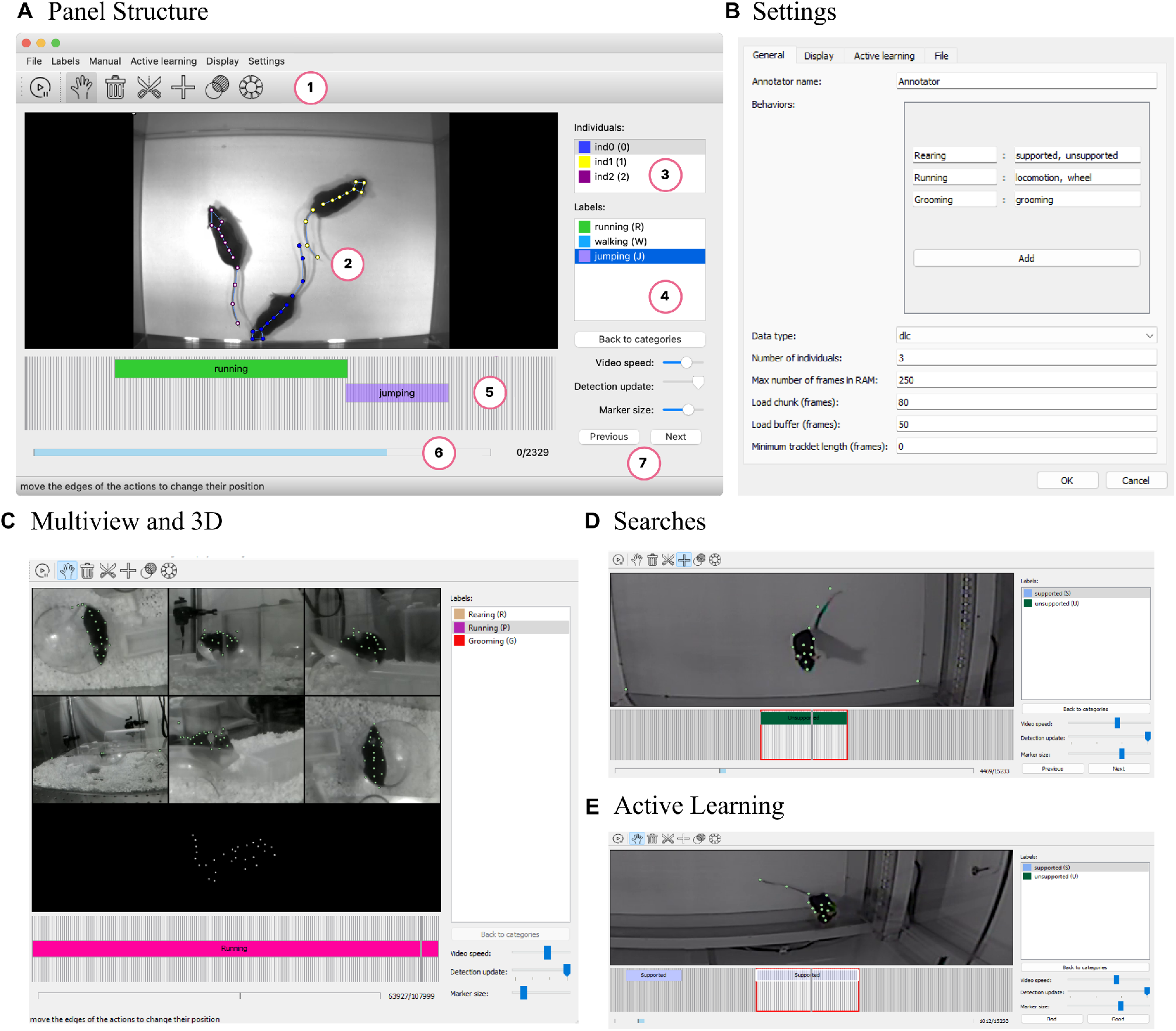
Graphic user interface for behavior annotations. (A) Panel structure. 1.Mode toolbar: includes Play/Pause, Drag, Delete, Cut, Add, Ambiguity, Switch actions. 2. Video canvas: Drag and zoom are supported, 3.Individuals menu 4.Labels menu: configuration of shortcuts and nested behaviors are supported. 5. Action bar: display annotated actions and has single-frame precision. 6. Global bar. Display current position in the video and loaded buffer. 7. Sliders: video speed slider (on pause mode can also increment by single frames), keypoints detection update and marker size, buttons to navigate in selected video set. One can annotate the behavior of multiple individuals; mouse image based on data from (24). (B) Settings. In the general settings, the user can select actions, type of input data, number of individual and depending on the machine capabilities the max number of frames in RAM, more settings are available for display, active learning and files. (C) Multiview and 3D. The GUI supports multiview in a mosaic setting for any number of selected videos. It also supports 3D visualization (bottom screen). Mouse images based on MausHaus (26, 57). (D) Searches. The annotation interface can guide the user (red window) towards given actions annotated or segments where no actions are annotated. (F) Active Learning. DLC2Action can provide action suggestions which can be visualized in the GUI (transparent action), after guidance toward the sample (red window) the user can subsequently validate or invalidate the s.uggestion. Mouse image in D/E based on data from (32)

The interface can be used for initial annotation, active learning, and the assessment or correction of automatic labels. The search feature of the annotation interface investigates video annotations and guides the user toward specific classes (Figure 6D). The data structures can also distinguish between negative and unseen frames. This is useful in the active learning scenario, where parts of the videos can be left unannotated. For an empirical assessment of prediction quality, the interface offers the option to sample a number of predicted intervals for each behavior class and manually grade them (Figure 6E).

### Customization and extensions

Because behavior annotation problems vary enormously in their structure — from discrete ethograms to hierarchical compositional labels, or from single-animal assays to multi-species camera traps — DLC2Action is designed for extensibility rather than coverage. Users can integrate new architectures, losses, metrics, and data formats by implementing a small number of standard interfaces without modifying the core training or evaluation pipeline. New annotation and keypoint data formats, data splitting methods, losses, metrics, models, feature extractors, and augmentations can be added. See the documentation at https://amathislab.github.io/DLC2action/html_docs/dlc2action.html for more information. To ensure stable performance, we also developed unit tests and employed continuous integration on GitHub. The code is tested on Windows, MacOS, and Ubuntu (Linux) systems.

## Discussion

DLC2Action demonstrates that frame-level behavior classification — long considered too annotation-intensive for broad adoption — is tractable across a wide range of organisms, assays, and recording modalities when temporal architectures are paired with systematic feature engineering and SSL training. The results across nine benchmarks establish a consistent pattern: temporal context is critical, pose-based kinematics sometimes suffice, and video or audio features add value precisely when the behavior of interest is not fully encoded in skeletal dynamics.

While numerous automated systems for accurate pose estimation have become available to expert biologists, allowing them to accurately study complex behaviors (20–26), manual behavior annotation is a laborious and time-consuming task. One can tackle this problem with unsupervised methods (10– 17, 19, 58), which require the validation of predicted clusters.

The architecture of DLC2Action is characterized by its compatibility with arbitrary multidimensional inputs, its inclusion of a specialized feature set for fine-grained behavioral phenotyping, and its modularity for incorporating state-of-the-art models. Furthermore, to address the need for standardized evaluation, we propose a formal benchmarking protocol on existing datasets to encourage the principled development and validation of subsequent models (Table 1). We found that DLC2action’s models outperform prior models across a wide range of settings (open field, social, and naturalistic behaviors). The near-saturation of 2D rodent action segmentation datasets likely reflects a combination of factors: there are only a few behaviors annotated, the kinematic signatures are relatively distinct, and the datasets are large enough to train temporal models to near-ceiling accuracy. Progress on these datasets will probably require either finer-grained behavioral ontologies. The 3D human datasets still have substantial room for improvement. This is particularly the case for naturalistic cooking, as presented in the EPFL-Smart-Kitchen-30 (43), and whole-body behavior, as presented in hBABEL (50, 54). These challenges stem from the inherent complexity of the annotated behavior, including its compositional and hierarchical nature. In summary, for some datasets, pose tracking alone is enough to achieve strong performance, while for others, significant gains can be achieved by combining video and pose modalities. Indeed, advances in recording technology are driving a trend towards multimodal datasets that integrate video, audio, IMU, and depth information (43, 46, 59). Our DLC2Action toolbox is designed to support this trend by providing a robust platform for annotating complex behaviors from these integrated data streams. DLC2Action complements other specialized tools focused on analyzing specific modalities, such as audio for vocalizations (60–62), or on jointly modeling behavioral and neural data (63).

A related practical consideration is the challenge of rare and imbalanced behaviors. In many behavioral studies, particularly in naturalistic settings, some behaviors occur infrequently (e.g. the EPFL-Smart-Kitchen-30 (43)), making their F1 scores difficult to interpret reliably. DLC2Action addresses this by providing users with class distribution reports at training initialization, showing the number of frames and behavioral bouts for each class across train/validation/test splits. This transparency allows to identify when rare classes are missing entirely from a split (which inflates performance estimates) or when test sets contain too few examples for confident predictions. We recommend that users flag classes with fewer than 10 test bouts as exploratory, examine per-class metrics rather than relying solely on macro-averaged F1 scores, and—when classes are severely under-represented—leverage DLC2Action’s active learning module to guide annotation efforts toward those classes. This guidance reflects a broader principle: behavior annotation is an iterative process, and users should understand the confidence limitations of their models before deploying them on new data. Future versions of DLC2Action could include automated warnings that alert users when rare classes lack sufficient test support, making these limitations more explicit and harder to overlook.

Currently, there is substantial advancement in comprehending behavior through the use of video-language models (43, 64–66). These models facilitate broad applications for various behavior understanding tasks (64) and could also be integrated into DLC2Action in the future.

To illustrate the capabilities of the toolbox, we also sought to identify robust links between eye movements and gameplay. We found that for some games, one could find such “strategies”. We note that we did not consider the state of the game or the screen, and thus were merely scratching the surface of this application. These patterns could be further studied in the context of decision making (67, 68), action anticipation (69), and attention shifts (70).

Overall, DLC2Action is an easy-to-use Python toolbox that includes numerous state-of-the-art action segmentation models. DLC2Action offers exceptional flexibility due to its ability to load data from a wide variety of data types (2D, 3D, or deep features) and formats. In addition, DLC2Action comes with an annotation GUI that allows users to generate ground-truth behavior annotations or assess the quality of the models’ predictions. DLC2Action addresses the challenges of manual annotation by leveraging pose estimation data, video features from powerful models such as DINOv3 (27) and VideoMAE (29, 30)., and self-supervised learning (SSL) to achieve high performance even with limited labeled datasets. The toolbox is characterized by its modularity, flexibility, and compatibility with diverse input formats, enabling users to analyze single-animal and multi-animal behaviors across various contexts. DLC2Action supports kinematic feature extraction, efficient hyperparameter optimization, and standardized benchmarking protocols, ensuring robust and reproducible results. The complementary open-source annotation GUI enhances usability by facilitating ground-truth generation and active learning. Leveraging open-source capabilities, DLC2Action could also be integrated with agentic systems like AmadeusGPT (57), which would provide a natural language interface for the analysis of behavior.

## Acknowledgments

We are grateful to many people for their feedback, alpha-testing, suggestions, and contributions, in particular to Lucas Stoffl, Margaret Lane, Marouane Jaakik, Steffen Schneider, Jennifer Shan, and Mackenzie Weygandt Mathis. We thank Daphne Bavalier and Dimitri Ryczko for their helpful feed-back on an earlier version of the manuscript. This work was funded by the Swiss National Science Foundation (SNSF) through grants 310030_212516 and 320030-227871.

## Author contributions

E.K. wrote the initial code and the GUIs. A.B. revised the code and the GUIs. A.B. carried out all experiments and made the figures. A.M., A.B., and E.K. wrote the manuscript. A.M. supervised the project.

## Methods

### Description of DLC2Action

#### Worflow

Following a project structure inspired by DeepLab-Cut (22), the tool organizes feature files, experiment history, and trained models within a single project folder (Figure S1A). The starting point is behavior annotations: time-stamped records of when each target behavior begins and ends. These can be created directly in our GUI or imported from annotation tools such as BORIS (71) or SimBA (34). Once a model is trained, its predictions can be reviewed and corrected within the same interface. If performance falls short, the user can refine hyperparameters, collect more annotations, or rely on integrated active learning algorithms to efficiently surface the most informative examples to label. The best-performing model — or an ensemble — can then be deployed to annotate new videos at scale (Figure S1A). Beyond this core loop, DLC2Action exposes a range of customizable options. Users select which kinematic features to extract from skeleton dynamics — such as speed, acceleration, joint angles, or keypoint distances (Figure S1B) — tailoring the representation to their specific behavioral questions. Data splitting is equally flexible: video-level splits test how well a model generalizes to entirely new recording conditions, making them well-suited for large-scale datasets, while time-based IID splits preserve temporal context and are better adapted to fine-grained behavioral analyzes (Figure S1C). Hyperparameters can be set manually or tuned automatically via an Optuna-based search (72). Metadata tags allow different subsets of data — for instance, recordings from different annotators — to be processed independently. Crucially, every hyperparameter search, training run, and prediction is logged in an experiment history that enforces consistency, prevents redundant computation, and gives users a transparent record they can query, reproduce, or build upon.

#### Feature extraction

The typical input to DLC2Action models is a set of features computed from pose estimation data. The user can use any combination of the following kinematic features: keypoint speeds, keypoint accelerations, distances between pairs of keypoints, distances from keypoints to the center between two individuals, polygon areas, angles, angular velocities, egocentric coordinates, allocentric coordinates, and centroid coordinates. Those features are computed for single individuals to study individual behavior classification or for pairs of individuals to classify their interactions. In addition, it is possible to load external pre-computed features and either concatenate them with those generated by DLC2Action or use them on their own. Feature arrays are cut into overlapping segments and fed into action segmentation models. For larger datasets, feature computation can be time- and memory-consuming, so DLC2Action keeps track of the features that have already been computed and reuses them in experiments run with the same data parameters. The features can be normalized according to the training set statistics after the dataset is split into subsets. These kinematic features have been shown to provide a substantial increase in performance on most models in all mouse datasets (Figure S10). Using kinematic features as input to the model generally enhances performance across most models and datasets (Figure S10). Since the evaluated behaviors can be linked to these kinematic features—for example, social interactions may be better predicted using distances between individuals—it is recommended to carefully choose kinematic features based on the target behaviors.

#### Training loss

The default loss function is adapted from (73) and is defined as a weighted sum of cross-entropy and a mean squared error consistency loss. The standard cross-entropy loss can be replaced with a focal loss (74); by default, it is weighted in inverse proportion to class support in the dataset. It is summed with self-supervised task losses with variable coefficients if they are used in the experiment. Custom losses can easily be added and used instead of the default loss.

#### Network architectures

DLC2Action incorporates eight models that perform the best on RGB-based action segmentation tasks: MS-TCN++ (38), EDTCN (75), C2F-TCN (41) and ASFormer (40). These models were selected because they achieve strong performance on computer vision benchmarks such as 50 salads (76) and breakfast (77). We do note that these benchmarks are for video-based action segmentation, and one typically utilizes pre-extracted deep visual features from video CNNs (30). We adapted these architectures to (also) work directly on pose or features derived from pose estimation data. Finally, we added MotionBERT (42), a model optimized for pose-based action recognition, which provides good performance, particularly on 3D human datasets.

In addition, we developed three adapted architectures that have been optimized for pose estimation data: MS-TCN3, C2F-Transformer, and MP-Transformer (Figure S1D). The main difference between MS-TCN3 and MS-TCN++ is the combination of the outputs from the last and second-to-last layers of the first stage to be passed as input to the second stage, allowing for richer representations (as opposed to only taking the output of the last layer). C2F-Transformer is a modification of C2F-TCN that replaces some convolution operations with attention mechanisms. Finally, MP-Transformer is a modification of a simple Transformer-Encoder architecture with additional max-pooling and up-sampling layers. Training time varies across architectures (Figure S2D provides insight into their relative computational efficiency). Throughout the pipeline, DLC2Action considers that, in practice, a large portion of the input data is often unannotated. All methods are adapted for missing data, and all models are separated into feature extraction and prediction generation modules, allowing them to be stacked with SSL modules.

Currently, DLC2Action supports contrastive learning, contrastive regression (78), time cycle consistency (79), order prediction (80), reverse order classification, and masked feature prediction. Any combination of those tasks can be added to any experiment automatically. Alternatively, it is also possible to train a model on just the self-supervised tasks and export the resulting features for analysis. The performance of the models varies depending on the datasets; therefore, it is recommended to optimize and train several of them when starting with a new dataset and either choose a model that performs the best or average the predictions from each model (Figure 4). DLC2Action also provides a choice of augmentations that can be applied to data to enhance training. They currently include rotation, mirroring, shifting, zooming, reordering individuals for interaction classification, adding Gaussian noise, masking keypoints, and temporal sub-sampling.

#### Hyperparameter search

Model training is defined by many hyperparameters, including the parameters of the model, the loss, the augmentations, and the feature extraction. The best configuration can be found automatically with the help of auto-ML algorithms. In particular, DLC2Action uses the Optuna (72) package to sample the hyperparameter space efficiently and prune bad experiments. The parameters and results of hyperparameter searches are saved in the project memory and can be loaded to update the project settings.

The segment length (i.e., the size of the time windows provided to the models) is a critical parameter and should be adjusted to match the specific behaviors of interest. Consequently, behaviors that differ greatly in their temporal dynamics may necessitate training distinct models, and users should resample or synchronize all modalities to a shared temporal resolution before training.

#### Splitting for cross validation

Datasets can be split into training, validation, and test data in several ways: random video selection, random segment selection (if segments are not overlapping), time split (e.g., the first 60% of each video can be used for training, the second 20% for validation, and the last 20% for testing), random selection with class balancing, and splitting according to file or sequence name, folder structure, or a split file. After certain split parameters are used in an experiment for the first time, DLC2Action remembers the selection and loads it whenever it is used again.

Changing the split parameters can help understand the amount of data needed to reach a given performance. The models’ performance depends on several parameters, including the number of labels, the distribution of labels, the relationship between labels and kinematic data, or the size of the training set. The latter is a major concern for experimentalists and has a strong influence on the models’ performance (Figure S2A-B). In the Open Field Test (32) dataset and with the MS-TCN3 model, we chose a time split using a fixed 20% for testing and variable training sizes (with at least 10% for validation). DLC2Action allows us to set the position of the validation segment, and in this experiment, we tested 10 different validation splits.

### Evaluation

Training logs and experiment history are saved by DLC2Action and can be analyzed subsequently. This facilitates initialization with previously trained models and the continuation of model training after interruption. In addition, it is possible to run evaluations on the validation subsets, the test subsets, or on new data and to generate predictions, which can be visualized in the annotation interface.

DLC2Action includes different metric functions for assessing performance both during and after training. They include segment-level metrics: segmental F1 and F-beta score, precision and recall, mAP; and frame-level metrics: frame-wise F1 and F-beta score, precision and recall, accuracy, edit distance, precision-recall AUC.

Most metrics can be reported either for each class separately or aggregated over classes. Moreover, it is possible to provide a meta tag with information such as annotator ID and choose the aggregation method over those tags. DLC2Action also includes various functions for storing all experiment-specific information in an internal structure format, and for aggregating results across multiple experiments (e.g., computing the mean and standard deviation of derived metrics).

### Active learning

The required amount of annotation (labeling) can be unclear when using DLC2Action models. Because model performance depends strongly on the amount of labeled input data (Figure S2A-B), DLC2Action includes an active learning module that identifies the most informative unlabeled segments for annotation, allowing users to prioritize labeling efforts rather than annotating uniformly. For example, in the OFT dataset, there are 275 annotated segments of grooming, 3003 segments of supported rearing, and 2090 segments of unsupported rearing. We recommend that users consider the quantities in terms of the number of segments and classes to estimate the amount of manual annotation required.

After optimization and training are performed, if the results are not satisfactory, it is possible to collect more data using active learning methods. The general idea behind active learning is to utilize the model’s uncertainty. There are several methods for assessing that uncertainty on a sample, and the most widely used in the context of deep learning are Bayesian Active Learning by Disagreement (81), least confidence, and entropy (82) scores. All of them are integrated into DLC2Action. After frame-wise scores are generated, DLC2Action suggests intervals with the highest scores for annotation (see Algorithm 1).

Ultimately, the method will generate a file that can be opened with our annotation interface.

### Datasets and experimental details

Here we describe the specifications of the datasets used to evaluate the performance of DLC2Action models and demonstrate some of their features.

We performed feature extraction as described above. In case additional features were available (e.g., TREBA features for the CalMS21 challenge (45), or pre-computed descriptors for SIMBA datasets (34)), they were also loaded and concatenated with the others. This was done to have comparable results to the classifiers used in those publications.

The metric used for comparison is frame-wise F1 score, macro-averaged across classes (excluding background where relevant) and annotators.

#### Multimodal action segmentation

All action segmentation results were generated using DLC2Action, as detailed in (43).

##### Algorithm 1: Interval Selection Algorithm

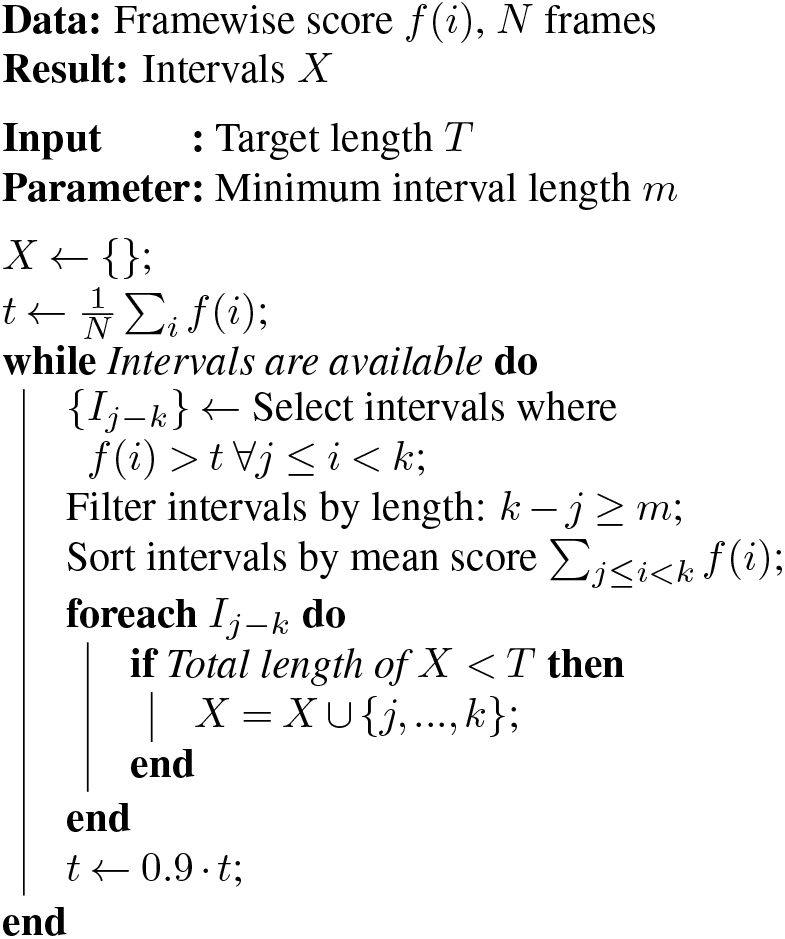

This reference also provides a comprehensive description of the VideoMAE pretraining on the ESK dataset and reports on model performance under other conditions. For the ESK dataset, we evaluate the C2F-TCN, MS-TCN3, EDTCN, and C2F-Transformer architectures. For the OFT dataset, we additionally include the MP-Transformer model. Notably, fine-tuning the VideoMAE model directly on the OFT dataset was unsuccessful, which we attribute to the limited amount of available training data.

For the MammAlps dataset (46), we used the detected bounding boxes and labeled bounding boxes as cropping boundaries for each individual before sending them as individual clips. We used the raw collected data to turn the dataset into an action segmentation dataset. Frames that do not have a detection for the given individuals were turned into black frames. The final dataset was composed of 953,456 labeled frames following a long-tail label distribution. We split the dataset randomly with a 70%/30% train/test split.

#### Single mouse with 2D pose: OFT and EPM datasets

We tested DLC2Action on two datasets presented by Sturman et al. (32). They comprise widely used behavioral tests in rodents: Open Field Test (OFT) and Elevated Plus Maze (EPM) and contain non-interactive behaviors annotated in a single-label fashion. The classes are supported rearing, unsupported rearing, and grooming for OFT and protected stretch, unprotected stretch, rearing, and grooming for EPM. The datasets consist of 10-minute videos, overall 20 for the OFT test and five for EPM. We followed the same leave-one-out split referenced in the publication in our experiments for the OFT dataset. For splitting the data of the EPM dataset into train and test subsets, we used a five-fold cross-validation strategy. The splits were generated by choosing the n-th 20% of consecutive frames in each video as the validation set for the n-th split (i.e., the first 5th, the second 5th, etc.). The results are aggregated over these splits.

#### Multiple rodents with 2D pose: CalMS21

In this work, we only consider CalMS21 task-1, from which the train/test split was defined in the challenge (45). Each CalMS21 experiment was repeated five times on the same data to assess consistency. The standard training procedure with only the annotated data was used.

#### Multiple rodents with 2D pose: SimBA datasets

We applied DLC2Action to datasets analyzed in (34), namely CRIM13 (Caltech Resident-Intruder Mouse 13) (47) and the rat resident-intruder dataset (SimBA-RAT) introduced in their paper. They both contain multi-label annotation of mostly social behaviors. The mouse dataset has eleven labels: approach, attack, chase, circle, copulation, drink, eat, sniff, up, clean, and walk away – and the rat dataset has seven: attack, anogenital sniffing, lateral threat, approach, boxing, avoidance, and submission. We used the features, annotations, and pose estimation data published in the SIMBA OSF repository (136k frames for the rat data and 842k frames for CRIM13) and compared DLC2Action models to a random forest classifier with SMOTE (default model in SimBA (52)). We used the same time-based split as previously described for the EPM dataset. Note that the results reported in (34) on CRIM13 and SimBA-RAT used a random frame-level data splitting method. Since this approach overestimates model performance and poorly reflects real-world generalization capabilities (to novel sessions from the same experiments), we made a new split for the community.

#### Single human with 3D pose: Shot7M2 and Hbabel

Finally, we tested DLC2Action in two 3D human pose estimation and hierarchical action segmentation datasets, namely Shot7M2 and hBABEL (19). In both datasets, we used the official splits for training and testing. Standard deviations are calculated over ten different model initializations. Because MotionBERT (42) takes a relatively long time to train, we tested two different model initializations. For conciseness in hBABEL, we report the F1 scores of the top 60 most frequent actions in each hierarchical level.

#### Self-supervised learning

We introduced SSL modules that are commonly leveraged in the machine learning (63, 83, 84). SSL tasks are represented in DLC2Action by a class with four attributes: loss function, neural network module, data transformation, and type, the latter determining how data is passed between modules during training (Figure S12). In particular, SSL target tasks focus on reconstructing the input data, while regularization tasks compare features extracted from the main models with those extracted from the SSL module, providing robustness in the feature space against perturbations and domain shifts (85). Finally, consistency tasks compare the features extracted in a first step with those extracted in a second step; this can provide better generalization and performance on downstream tasks (86). For the experiment (Figure S12D), we compared the C2F-TCN model when using an SSL Consistency module and a contrastive loss. We ran the model on five random sets of videos and used the remaining videos as part of the SSL training set and as part of the testing set.

#### ATARI-Head dataset

The ATARI-Head (53) dataset is a large-scale public dataset that combines eye gaze with controller commands from 20 subjects playing various ATARI games. Each game was independently assessed using the MS-TCN3, C2F-TCN, MP-Transformer, and MLP models. Chance level was calculated using the following formula:

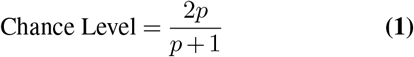

Where *p* is the marginal probability *p* = *p*(*action* = *True*) for a specific game. The dataset was split into 80/20% train/test splits using entire videos (i.e. 304/96 total number of videos). The experiments were repeated with 3 different seeds for cross-validation. As the eye tracker had a higher sampling rate, we averaged per video frame (30 Hz).

#### Annotation Interface

The annotation interface allows users to manually annotate behaviors in videos using estimated 2D and 3D pose. Below we provide an exhaustive list of the available features.

### Project management

The first window creates and loads DLC2Action annotation projects. All annotations and data references are saved in project-specific folders.

### Main annotation interface

The main interface provides a visualization of the videos of interest (video panel), a frame-accurate navigation bar showing annotated behaviors, a video-level navigation bar, a list of labels with associated colors and shortcuts, and a toolbar for editing annotations.

### Toolbar

The toolbar includes a displacement tool, a delete tool, an add tool, and a confusion tool (which assigns an uncertainty score to an annotation).

### Labels

Labels can be edited and organized in a two-level nested structure, enabling annotation of many behaviors with a limited number of shortcuts.

### Display

The video panel can display one or multiple videos simultaneously, together with 2D pose estimation data from DeepLabCut (20).

### Searches

The interface provides several search options to help identify missing annotations. Users can search for segments that are unlabeled or labeled with a specific behavior.

### Active learning

Integrating the annotation interface with DLC2Action enables active model optimization. The interface can load DLC2Action model predictions, and users can provide feedback (correct/incorrect), focusing annotation effort on segments where behaviors are most likely to occur.

### Settings

Additional settings in the settings panel allow users to manage the memory buffer and to adjust display and active learning parameters.

## Supplementary Figures

### Workflow and models

**Figure S1.**
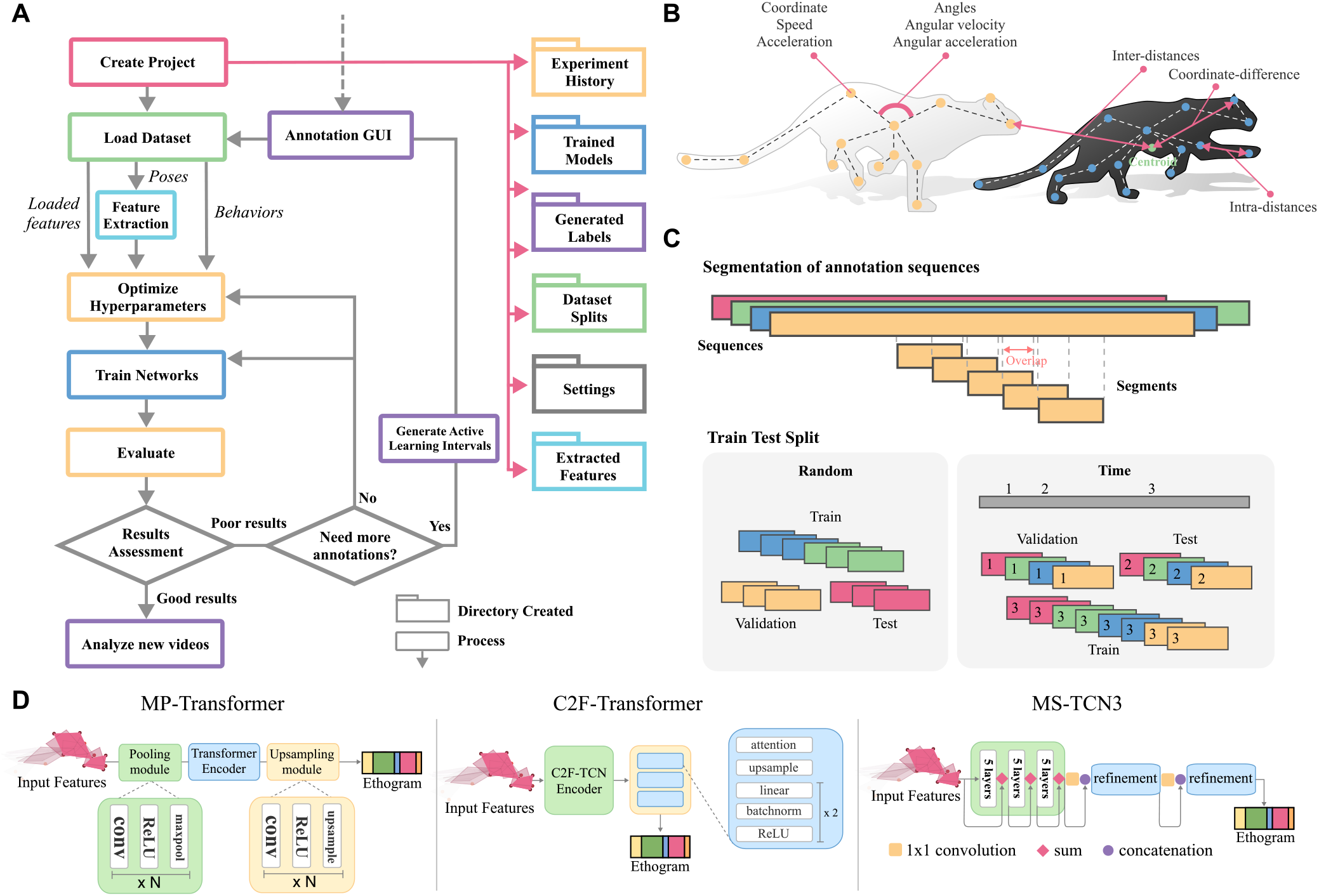
Worflow, feature extraction and preprocessing. (A) Workflow. DLC2Action covers every step of the behavior annotation workflow using a supervised learning approach. Specifically, projects are created to store the weights of trained models and dataset information. After data loading, the user can automatically optimize models’ hyperparameters (configurable part of the models’ learning process) and train neural network on their action segmentation dataset. The evaluation of the models gives cues on whether to annotate more data and active learning can be used to guide the annotation process using the provided GUI. Finally the user can analyze new videos using the trained models. (B) Feature extraction DLC2Action can extract the position, speed, acceleration of each keypoint, the angles, angular velocity and angular acceleration of each joints, the pair-wise distances between keypoints for specific individuals and between individuals. (C) Segmentation and Splitting methods. All annotation sequences can be divided in overlapping segments. Considering a training, testing and validation set, the “random” method sort sequences into subsets randomly and the “time” method splits each sequence into training, validation and test subsequences. Other splitting methods are available in DLC2Action (see Methods). (D) Model architecture changes in DLC2Action. All models were changed to accept kinematic (or multimodal) input instead of video. (Left) MP-Transformer includes additional max-pooling and upsampling with respect to a standard transformer-encoder architecture. (Middle) C2F-Transformer replaces convolution operations with attention layers from the C2F-TCN architecture (41). (Right) MS-TCN3 combines the output of the last and second-to-last layers of the first stage as input to the second stage. Adapted from MS-TCN++ (38).

### Size and performance

**Figure S2.**
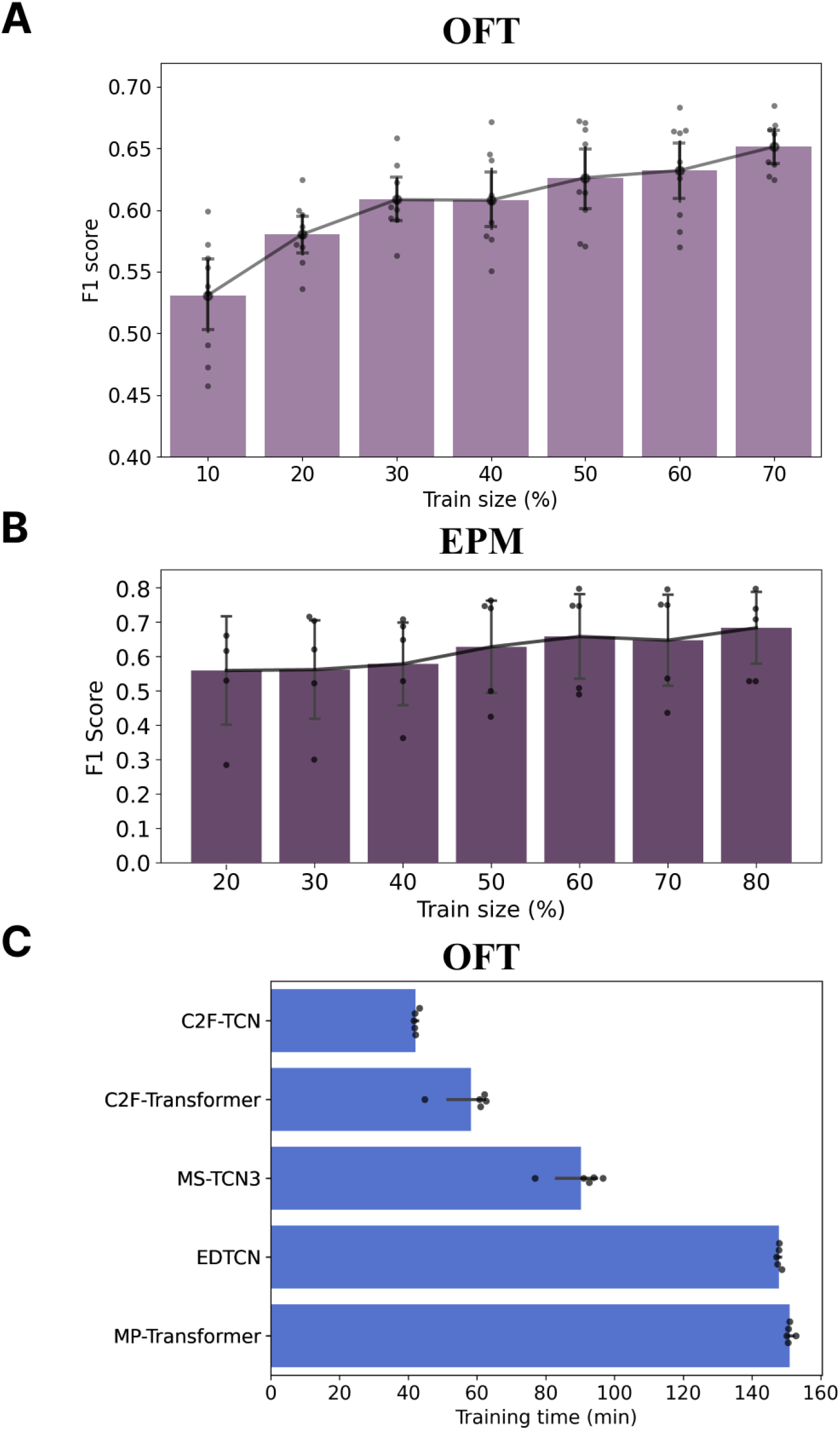
Effect of size of the training set and Training time. (A) F1-score against the proportion of the dataset used for training, evaluation was done on one video according to a leave-one-out split. Experimented on the OFT dataset (32) with the MS-TCN3 model (m=20 splits). (B) F1-score against the proportion of the dataset used for training, evaluation was done on 20% of the videos according to a time:strict split. Experimented on the EPM dataset (32) with the C2F-TCN model (m=5 splits). (C) Duration of training for 100 epochs. Experimented on the OFT dataset (32) for four different models with a batch size of 256, a segment length of 256 frames with 200 frames overlap, kinematic features, and using a GeForce RTX 3090 (m=5 splits).

### Detailed benchmark results

**Figure S3.**
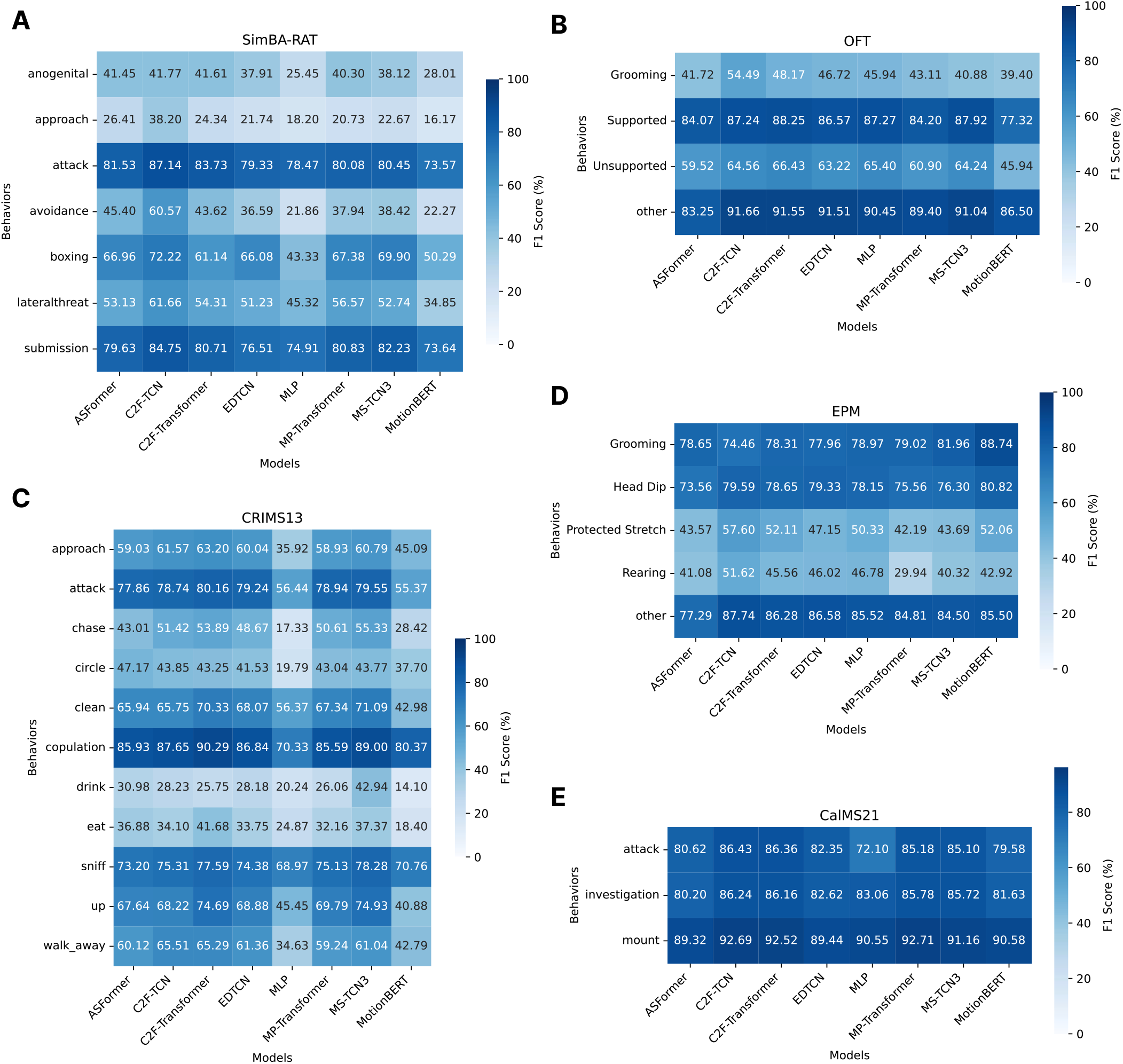
Average F1-scores per class for rodent datasets Average over different splits (n) or initialization seeds for each model (m). (A) Results for SimBA-RAT (34) (m=5) (B) Results for Open Field Test (32) (m=20) (C) Results for CRIMS13 (47) (m=5) (D) Results for Elevated platform maze (32) (m=5) (E) Results for CalMS21 (45) (n=5)

**Figure S4.**
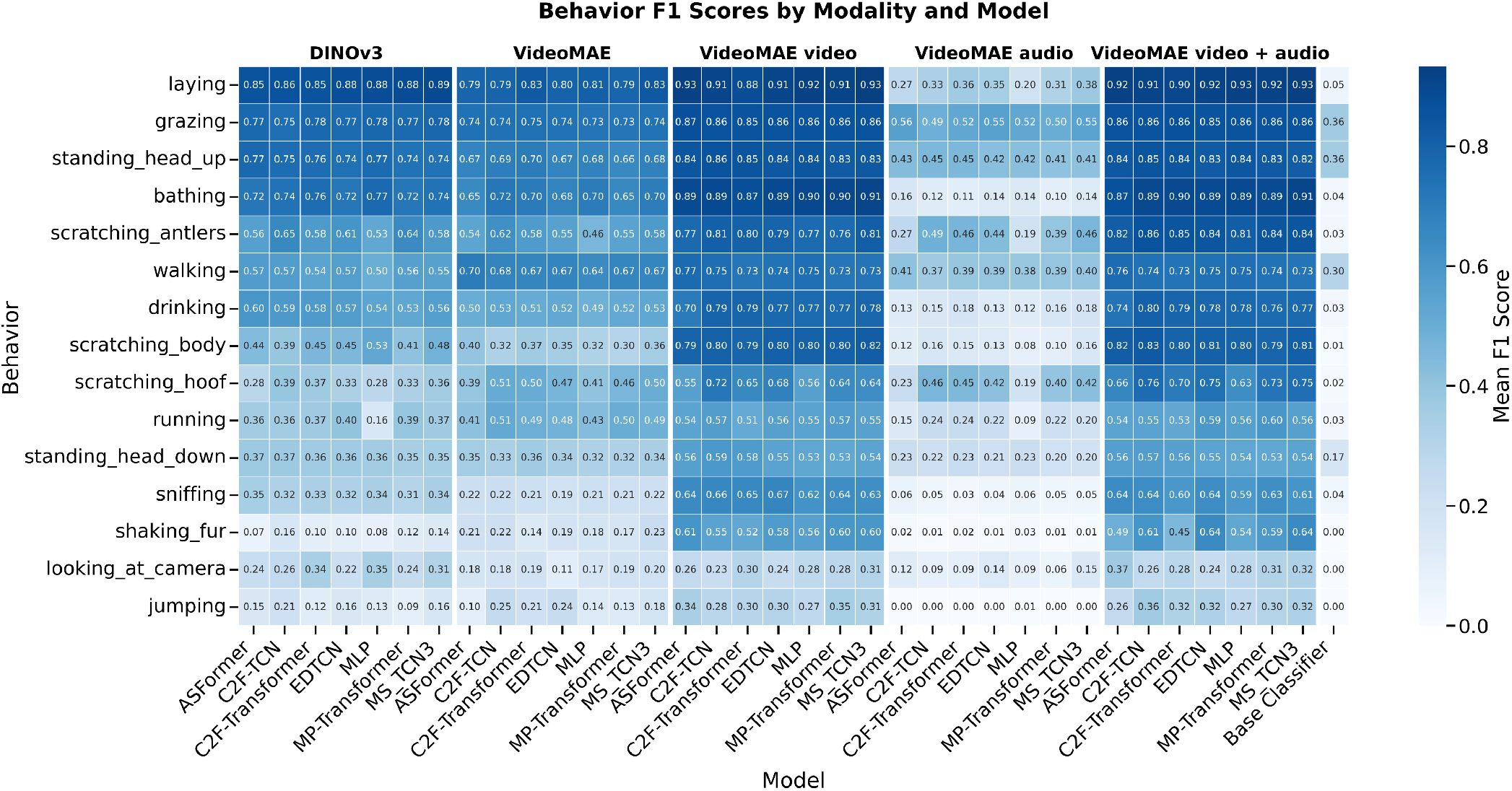
Average F1-scores per class and per modality in the MammAlps dataset (46). We differ video embeddings extracted by DINOv3 (27), VideoMAE pretrained on Kinetics700, and VideoMAE finetuned on MammAlps when using only the video (VideoMAE video), only the audio (VideoMAE audio), or both (VideoMAE video + audio.

**Figure S5.**
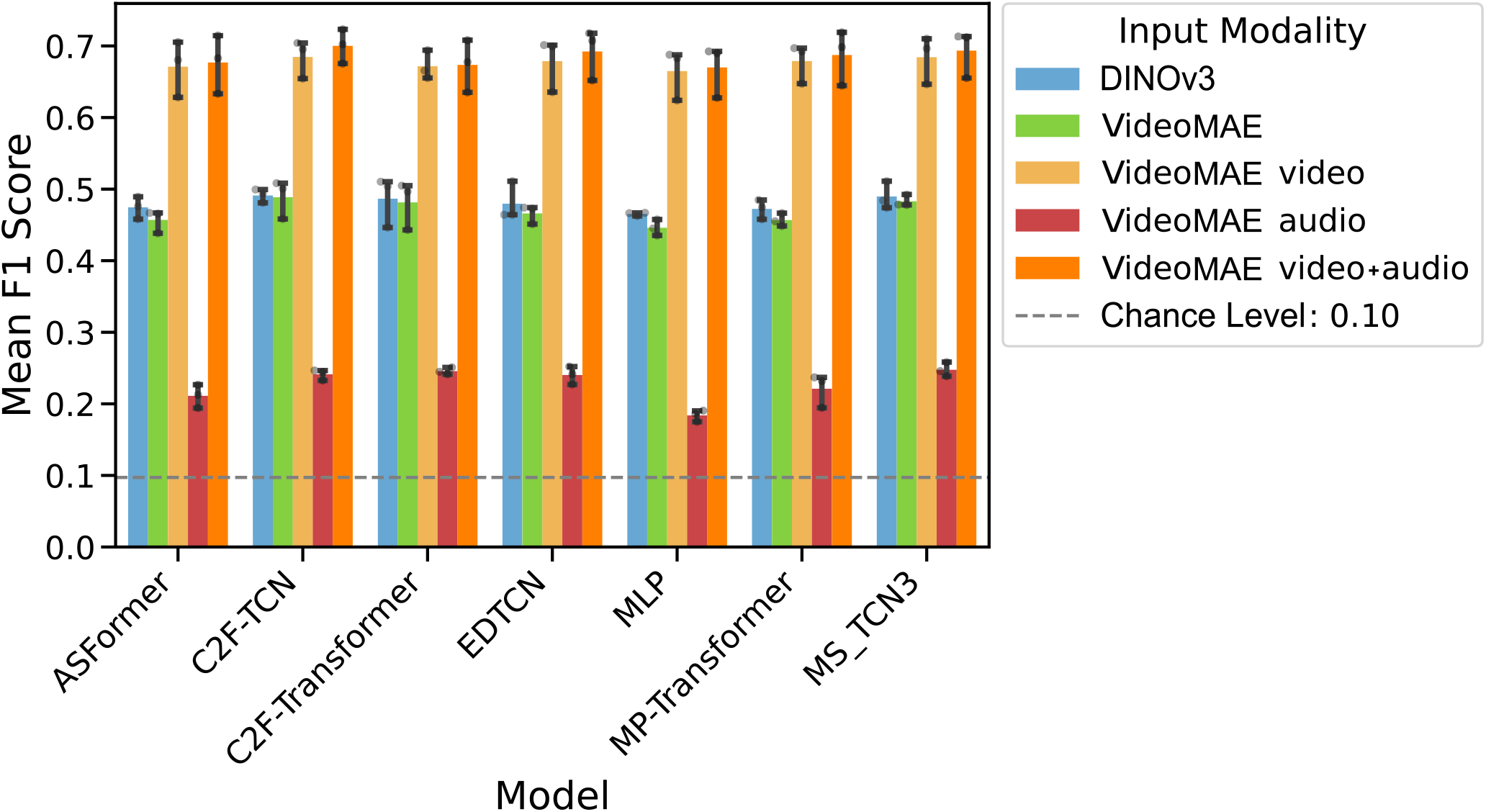
Average F1 score per model and modality. We differ video embeddings extracted by DINOv3 (27), VideoMAE pretrained on Kinetics700, and VideoMAE finetuned on MammAlps when using only the video (VideoMAE video), only the audio (VideoMAE audio), or both (VideoMAE video + audio.

### Multimodality gain in MammAlps

**Figure S6.**
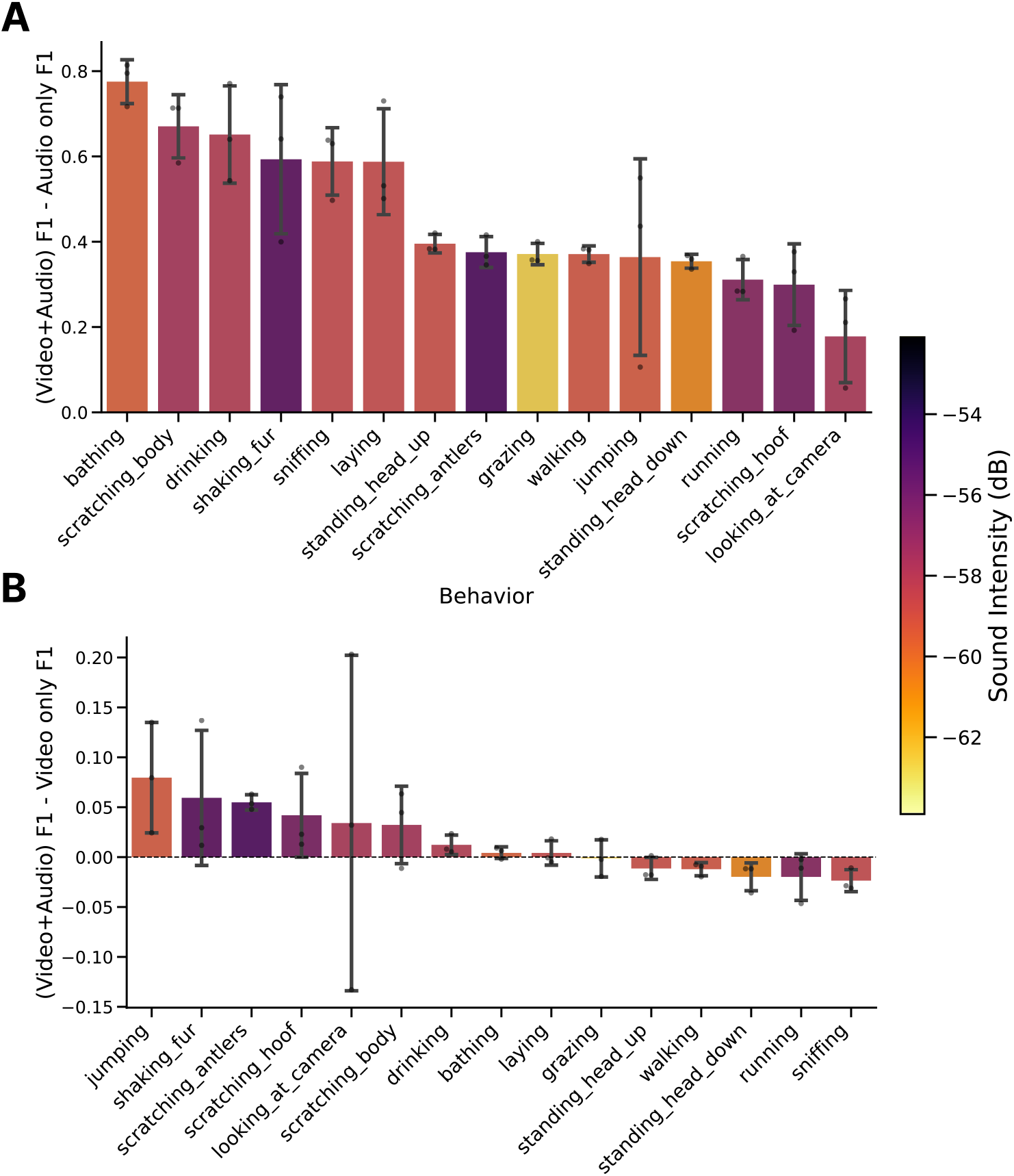
Performance gain of adding video to audio or audio to video per behavior in the MammAlps dataset. (A) Difference in F1 scores (Video + Audio) vs. Audio-only for each behavior. (B) Difference in F1 scores (Video + Audio) vs. Video-only for each behavior. Positive values indicate an improvement when adding audio (A) or video (B).

**Figure S7.**
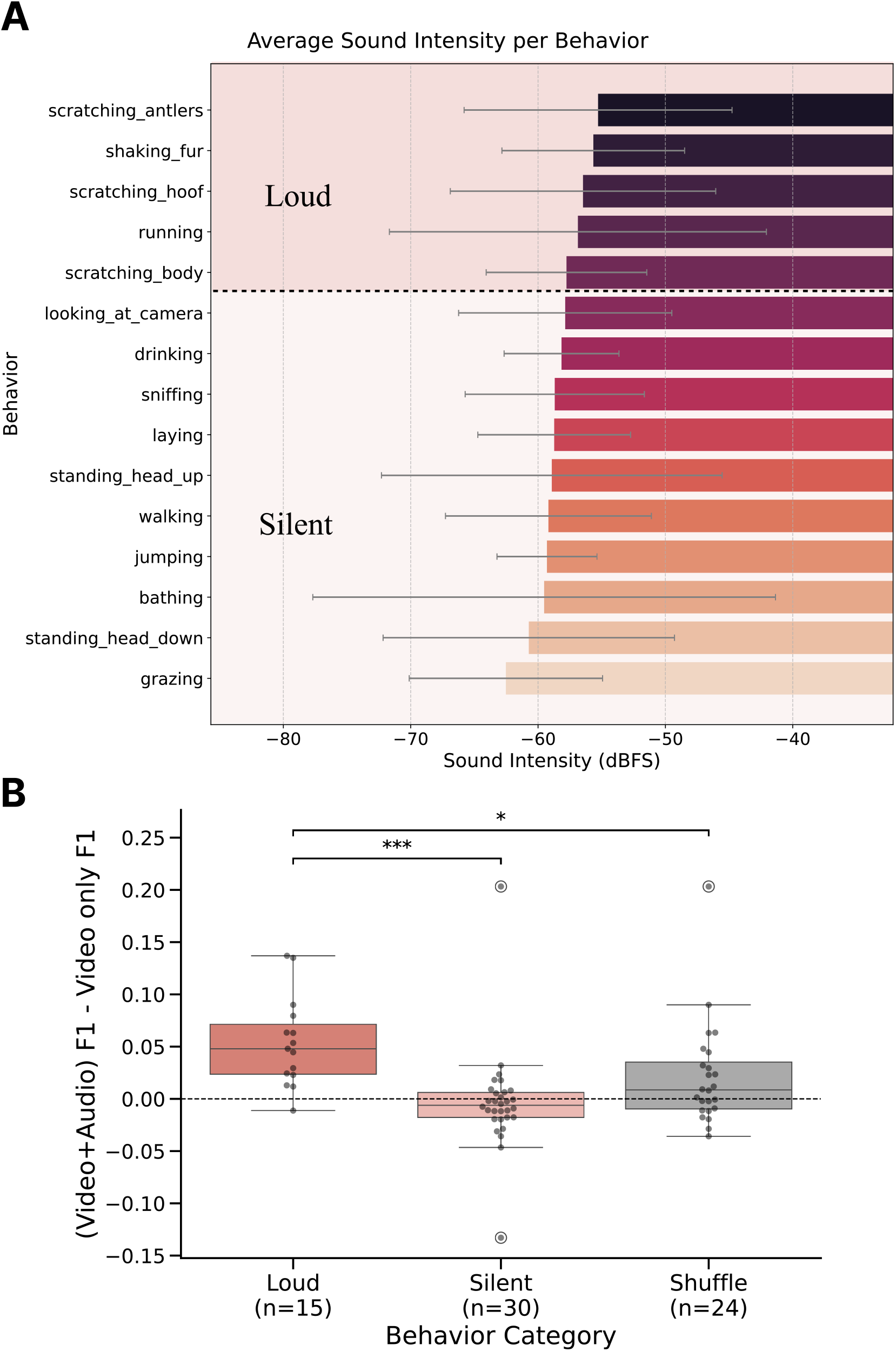
Sound intensity (dBFS) per behavior in the MammAlps dataset (46). (A) Error bars represent standard deviation across all recorded sequences. An arbitrary sound intensity threshold has been selected to separate “loud” behaviors from “silent” behaviors. (B) Effect of Audio modality on performance. Adding the audio modality has a significant effect on the classification of “loud” behaviors as defined in A. We divide the 15 behaviors into five loud behaviors and 10 silent behaviors based on their sound intensity (see FigureS7), across three different splits. Shuffle condition corresponds to the performance of a random subset of behaviors, done for each split, of eight behavioral labels irrespectively of their sound intensity. Error bars correspond to standard deviation over different model architectures (n). Statistical significance was determined using a Kruskal-Wallis with a post-hoc pairwise Mann-Whitney U (non-normal distribution). Asterisks denote statistically significant differences between the indicated groups (\**p <* 0.05, \*\**p <* 0.01, \*\*\**p <* 0.001). Comparisons unmarked were not statistically significant (*p >* 0.05).

### Splitting distribution in MammAlps

**Figure S8.**
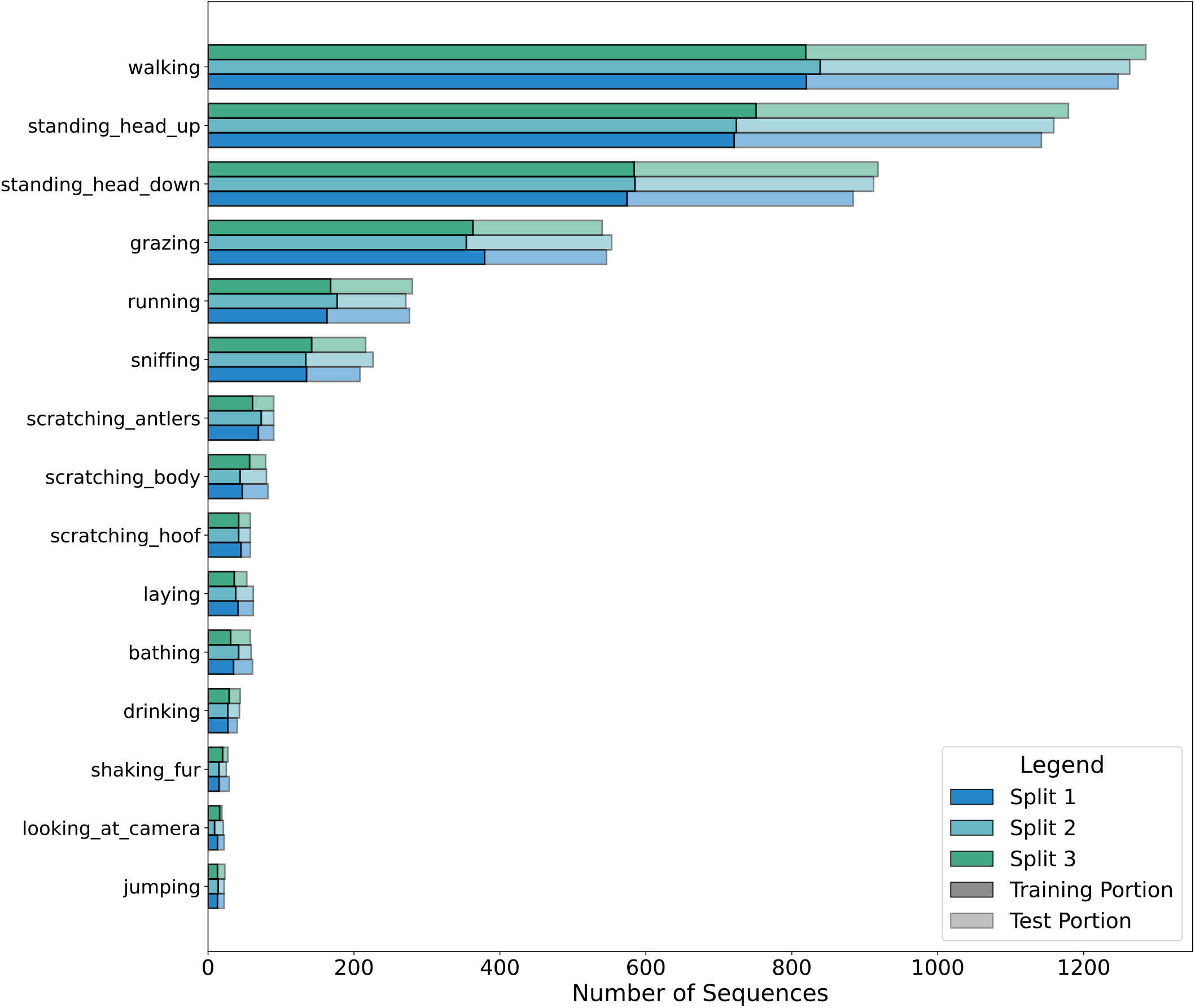
Distribution of behavior per split. Number of segment for each behavior in the train and test splits.

### Visual features on OFT

**Figure S9.**
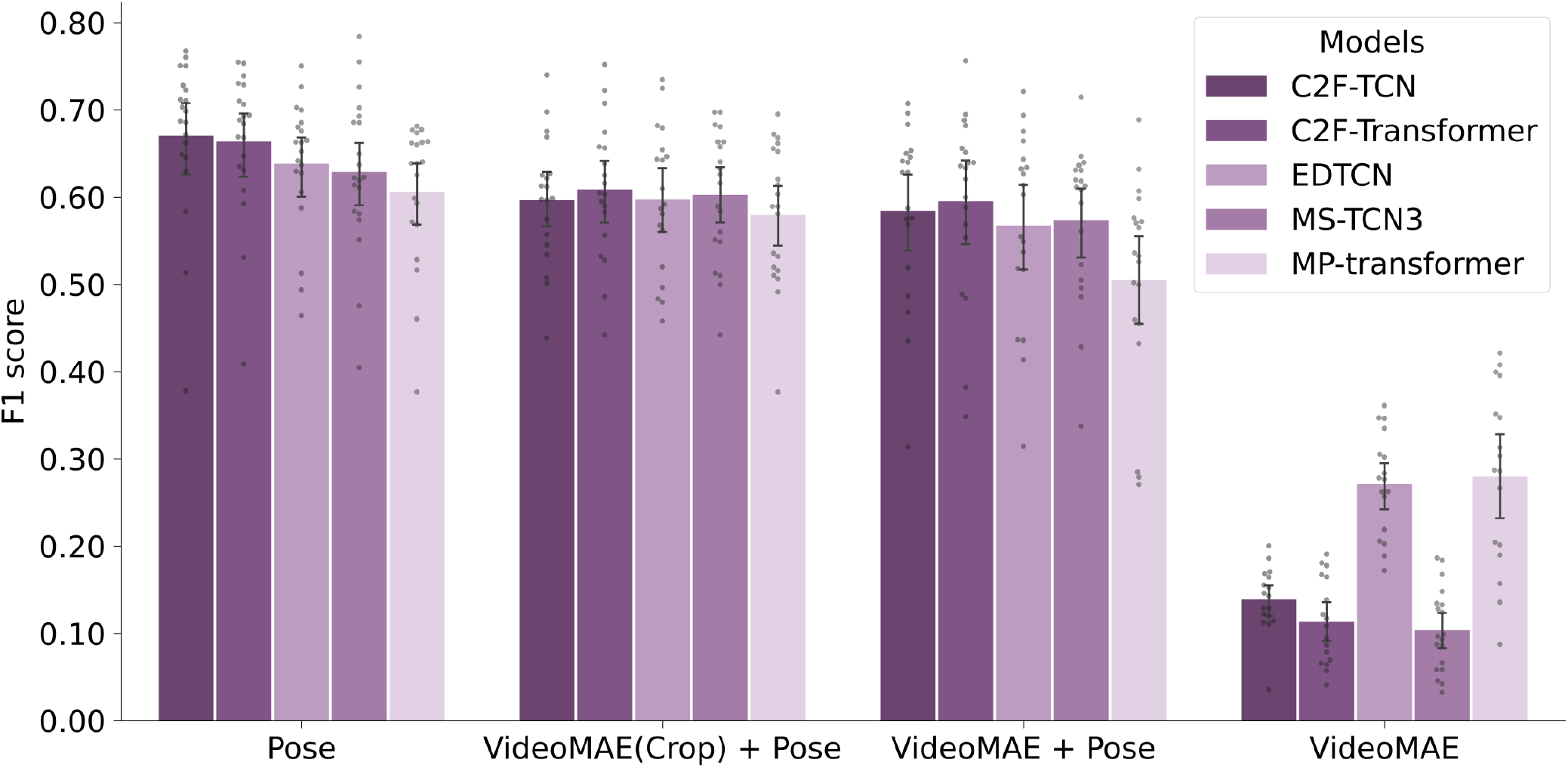
Performance of pre-trained visual features on the OFT dataset. Average F1-score of DLC2Action models trained with pose estimation input, pose estimation concatenated with VideoMAE features from images cropped around the mouse, pose estimation concatenated with VideoMAE features from original videos, VideoMAE features from original videos. Video frames are resized to 224×224 before tokenization. (n=20 splits)

### Importance of kinematic features

**Figure S10.**
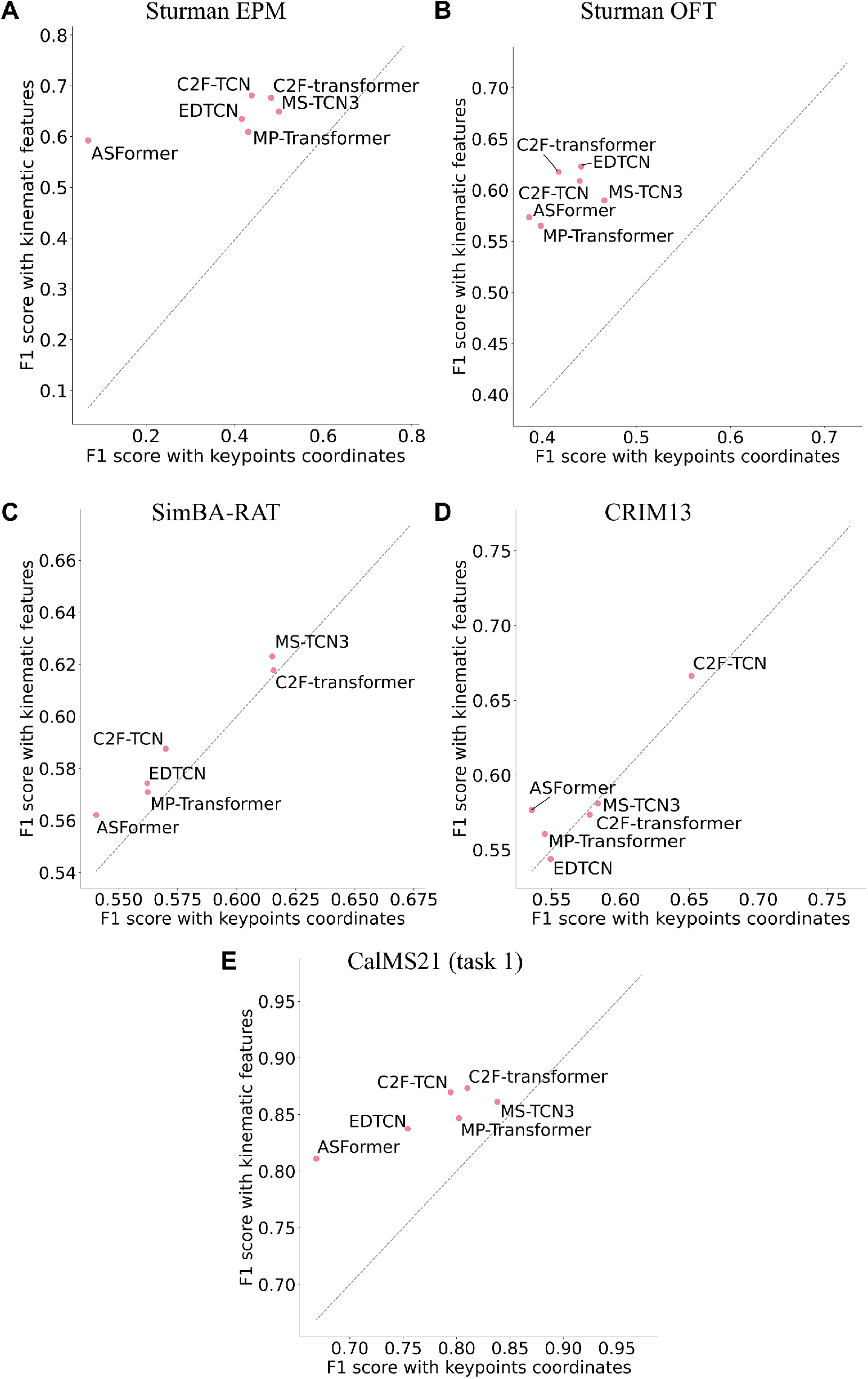
Comparison with/without kinematic features. For all 2D datasets and in most models, extracted kinematic features increases performances. Average F1 score over different splits (n) or model initialization seeds for each model (m) (A) Elevated platform maze (32) (EPM, n=5) (B) Open Field Test (32) (OFT, n=20) (C) SimBA RAT (34) (n=5) (D) CRIM13 (47) (n=5) (E) CalMS21 task 1 (45) (m=5)

### Statistical characteristics of DLC2Action predictions

**Figure S11.**
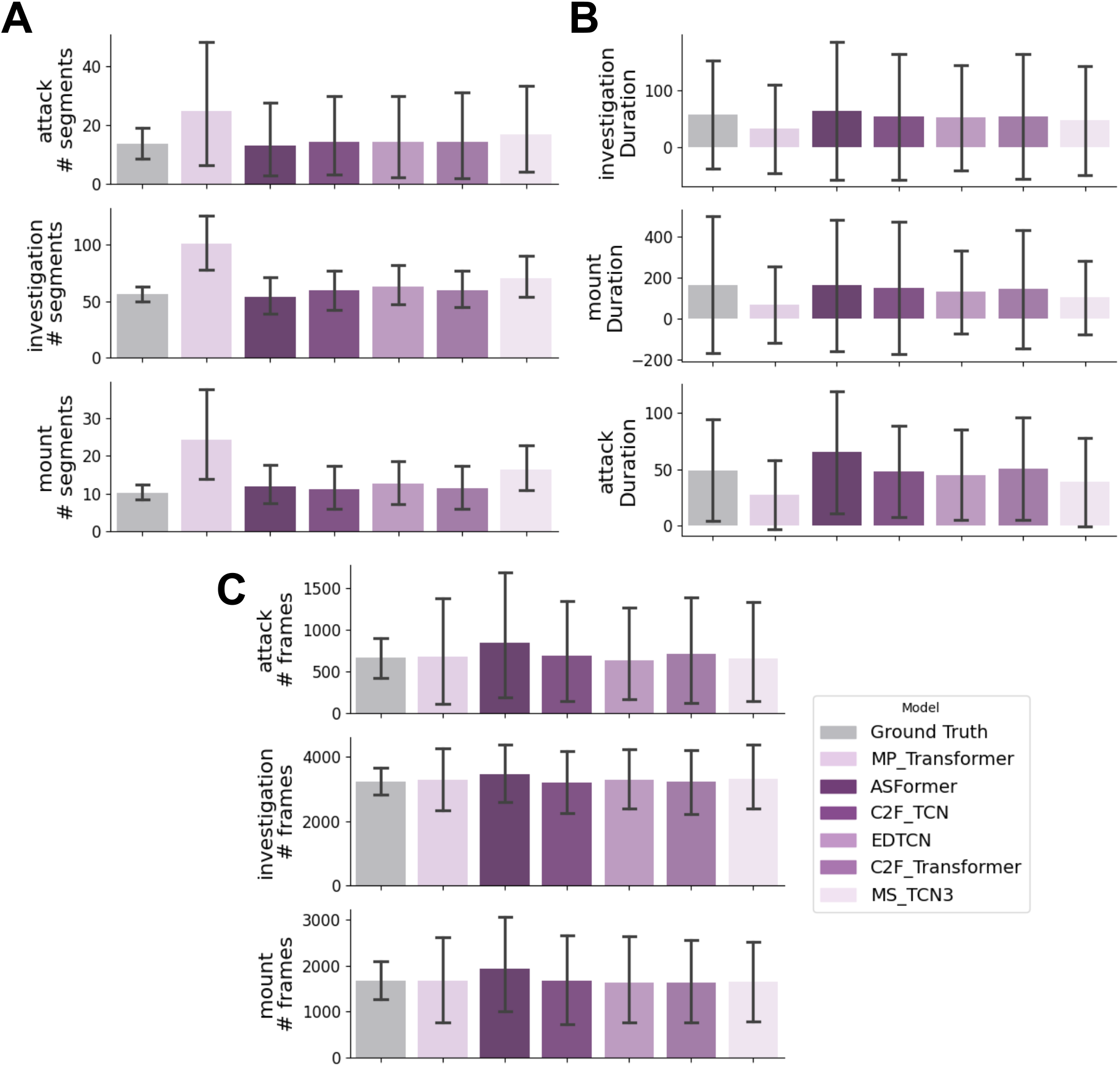
Statistical characteristics of DLC2Action model predictions for the CalMS21 dataset (45). (A) Number of segments. (B) Average duration of segments predicted. (C) Number of frames labeled

### Self-Supervised Learning

**Figure S12.**
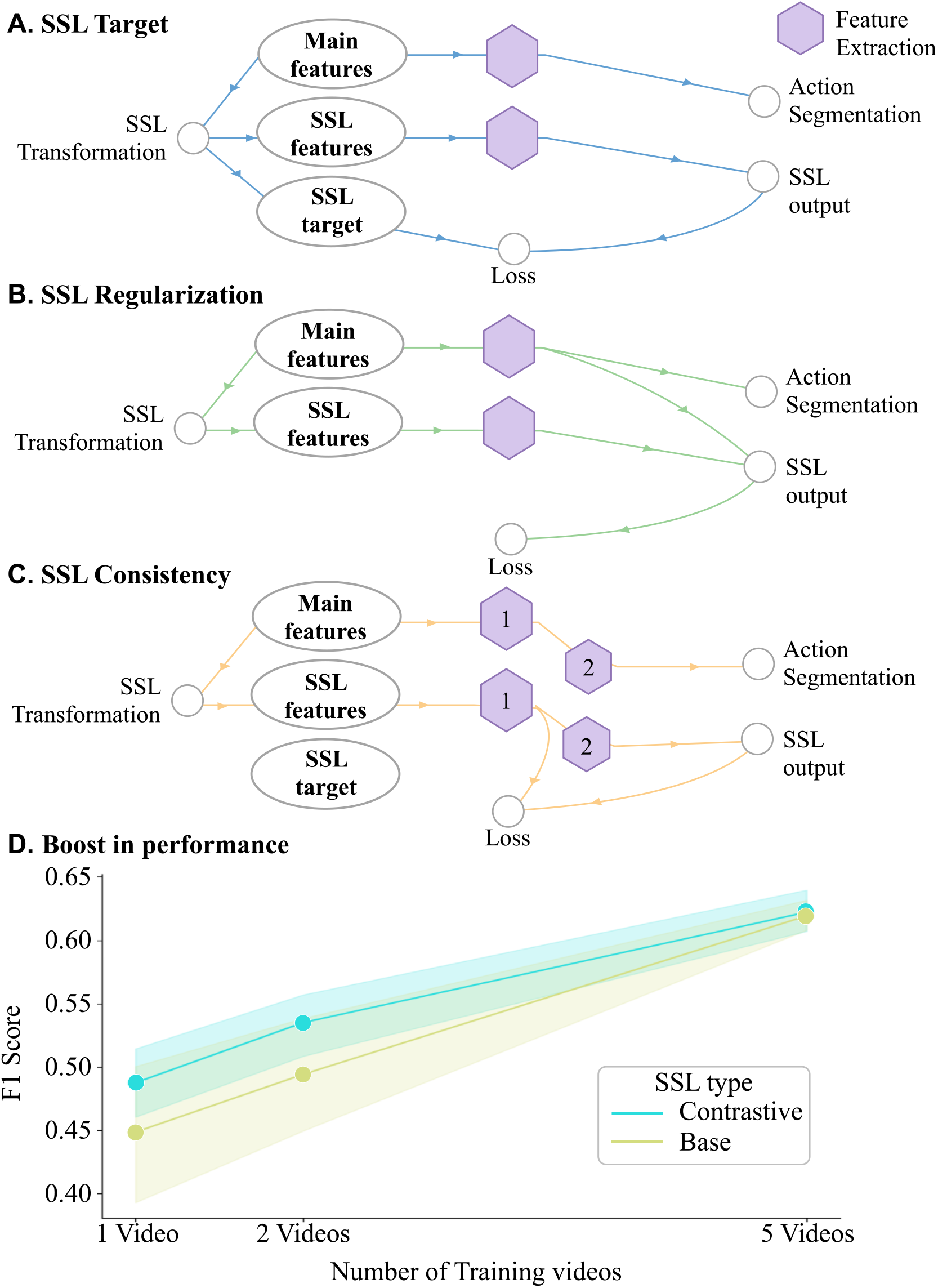
Diagram of supported SSL mechanisms and Results. SSL is a representation learning approach that learns feature representations from unlabeled data by exploiting the structure of the (input) data itself, providing models with richer, more informative representations that improve performance on downstream supervised tasks. Supported SSL types can be divided into three categories (A-C). (A) SSL target loss involves solving an input reconstruction task, comparing the SSL output to a pre-defined target, e.g. input data or kinematic features, to compute the loss. (B) SSL Regularization tasks compare the features extracted from the SSL module with those extracted during the action segmentation task to compute and update the loss. (C) Finally, SSL consistency tasks involve a two-step feature extraction process, comparing the features extracted in the first step with those extracted in the second step to update the loss. (D) Using unlabeled data can boost performance for small amount of labeled data. We report the SSL performance of an SSL module with a contrastive loss (one type of SSL Target) using 5% (1 video), 10% (2 videos) and 25% (5 videos) of labeled data and the whole OFT dataset (32) as part of the SSL training. (N = 5 independent experiments)

### Detailed results Atari-HEAD

**Figure S13.**
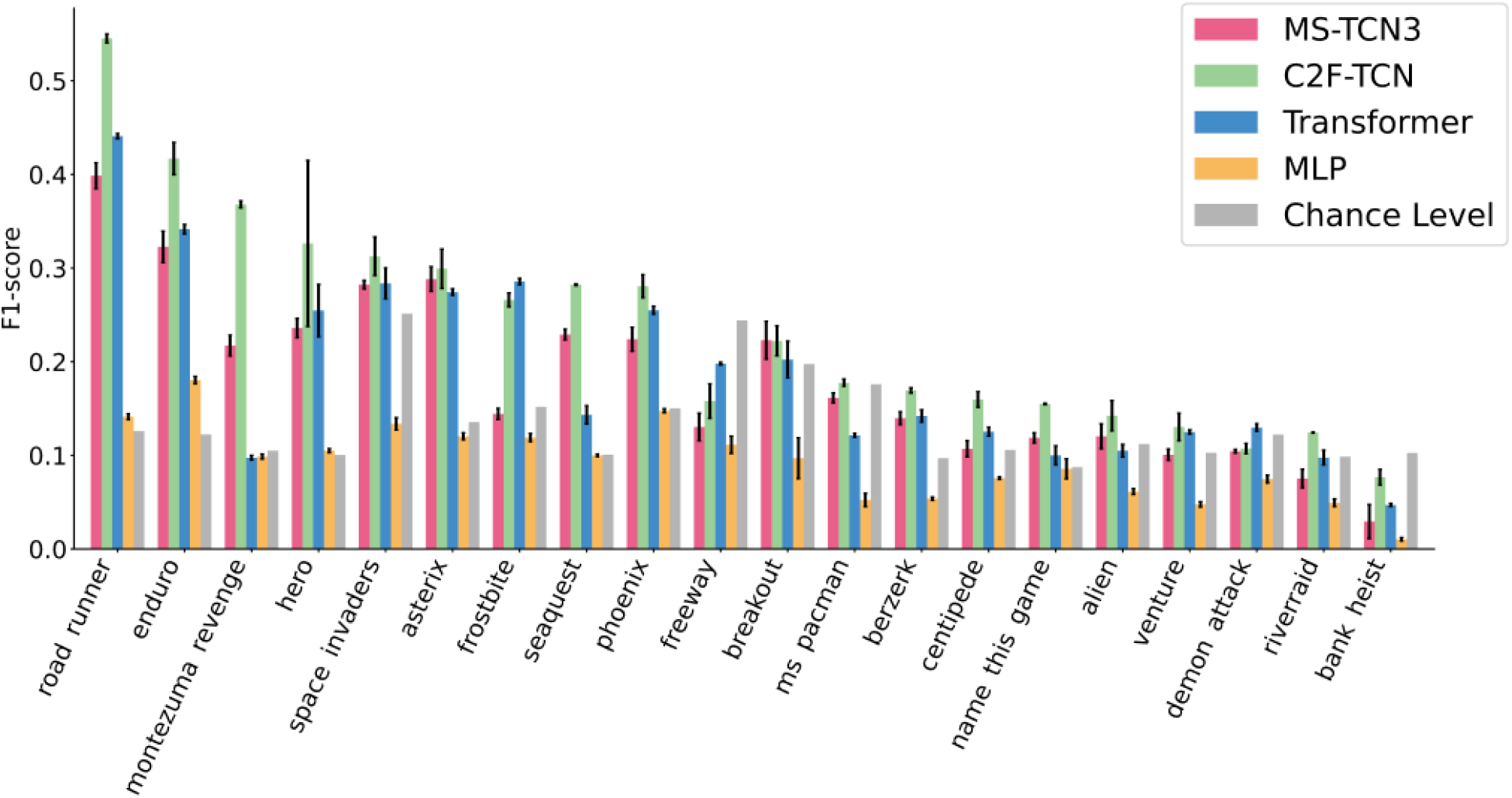
Performance of DLC2Action models on all 20 games in Atari-Head (53). We compare the MS-TCN3, the C2F-TCN, the MP-Transformer and the MLP model with chance level. F1 scores are averaged over game commands present in each game. Error bars correspond to the standard deviation over five data splits.

### Video examples

**Video 1.**
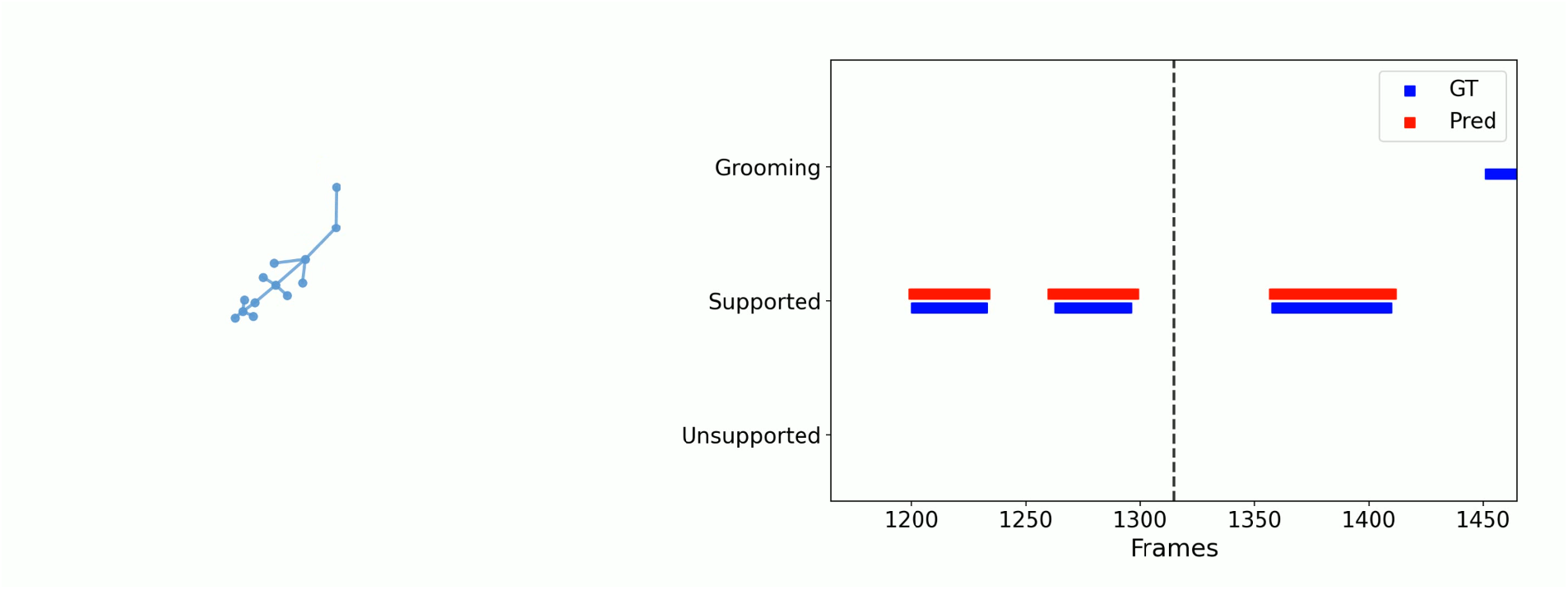
Here: A single frame of a video that illustrates DLC2Action on a single animal 2D pose dataset(OFT (32)). The attached video shows the input kinematics, the ground truth behavior annotations and the best DLC2Action model prediction (C2F-TCN).

**Video 2.**
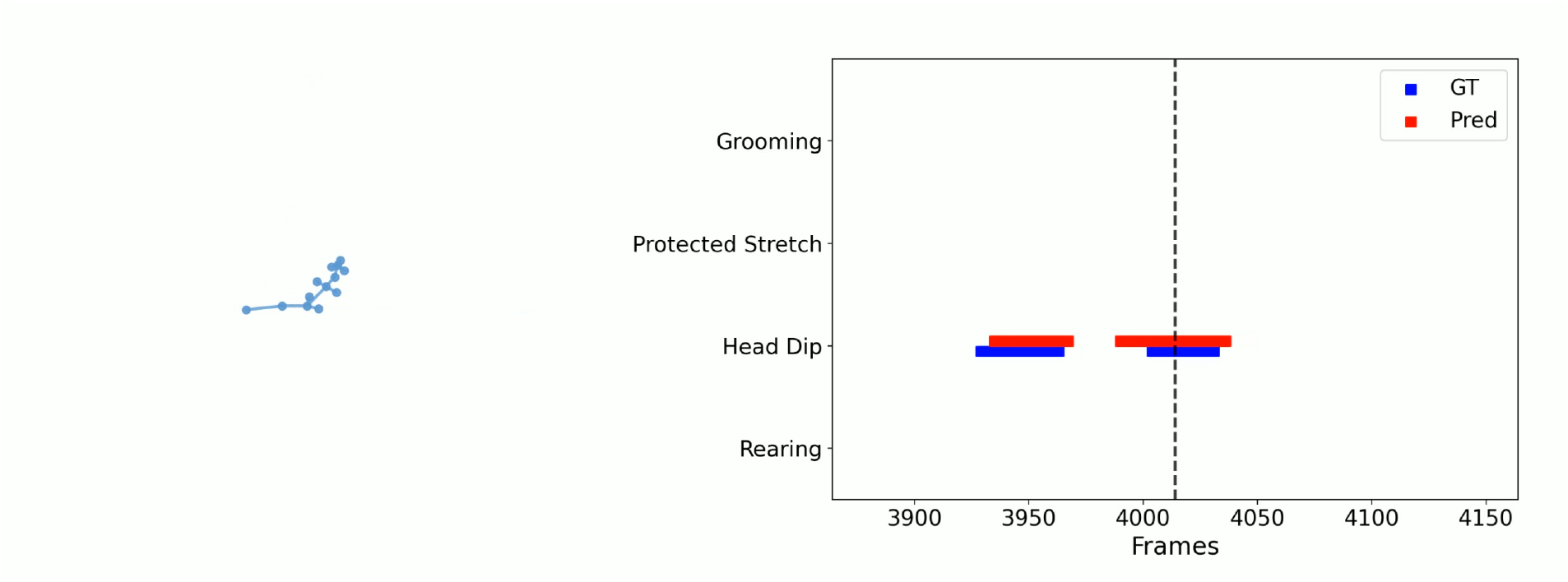
Here: A single frame of a video that illustrates DLC2Action on a single animal 2D pose dataset(EPM (32)). The attached video shows the input kinematics, the ground truth behavior annotations and the best DLC2Action model prediction (C2F-TCN).

**Video 3.**
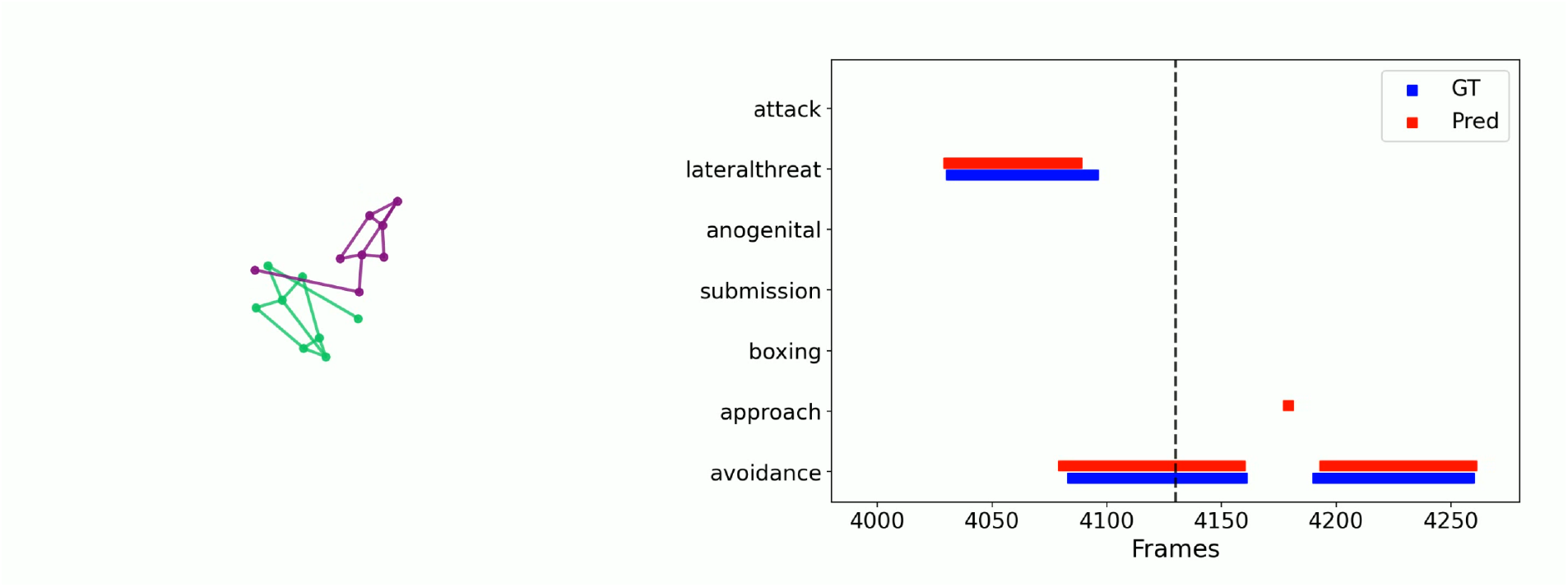
Here: A single frame of a video that illustrates DLC2Action on a multi-individual dataset (SimBA-RAT (34)). The attached video shows the input kinematics, the ground truth behavior annotations and the best DLC2Action model prediction (C2F-TCN).

**Video 4.**
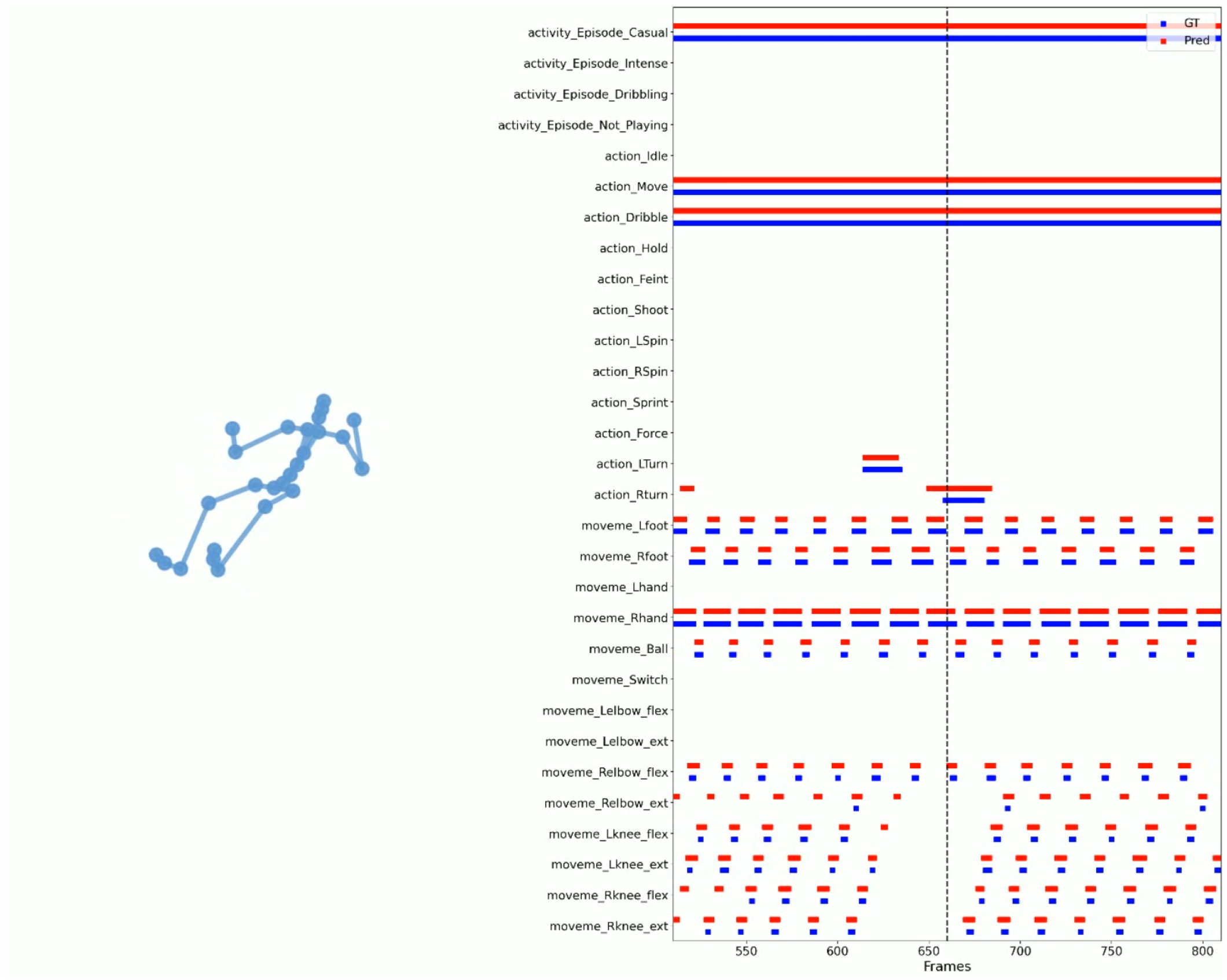
Here: A single frame of a video that illustrates DLC2Action on a 3D pose dataset(Shot7M2 (50)). The attached video shows the input kinematics, the ground truth behavior annotations and the best DLC2Action model prediction (ASFormer).

